# A stargate mechanism of *Microviridae* genome delivery unveiled by cryogenic electron tomography

**DOI:** 10.1101/2024.06.11.598214

**Authors:** Pavol Bardy, Conor I.W. MacDonald, Paul C. Kirchberger, Huw T. Jenkins, Tibor Botka, Lewis Byrom, Nawshin T.B. Alim, Daouda A.K. Traore, Hannah C. König, Tristan R. Nicholas, Maria Chechik, Samuel J. Hart, Johan P. Turkenburg, James N. Blaza, J. Thomas Beatty, Paul C.M. Fogg, Alfred A. Antson

## Abstract

Single-stranded DNA bacteriophages of the *Microviridae* family are major components of the global virosphere. Microviruses are highly abundant in aquatic ecosystems and are prominent members of the mammalian gut microbiome, where their diversity has been linked to various chronic health disorders. Despite the clear importance of microviruses, little is known about the molecular mechanism of host infection. Here, we have characterized an exceptionally large microvirus, Ebor, and provide crucial insights into long-standing mechanistic questions. Cryogenic electron microscopy of Ebor revealed a capsid with trimeric protrusions that recognise lipopolysaccharides on the host surface. Cryogenic electron tomography of the host cell colonized with virus particles demonstrated that the virus initially attaches to the cell via five such protrusions, located at the corners of a single pentamer. This interaction triggers a stargate mechanism of capsid opening along the 5-fold symmetry axis, enabling delivery of the virus genome. Despite variations in specific virus-host interactions among different *Microviridae* family viruses, structural data indicate that the stargate mechanism of infection is universally employed by all members of the family. Startlingly, our data reveal a mechanistic link for the opening of relatively small capsids made out of a single jelly-roll fold with the structurally unrelated giant viruses.

## Introduction

Viruses showcase extraordinary diversity in shape, symmetry, and size, with the genome sizes of giant viruses and microviruses varying by up to three orders of magnitude. In bacteriophages, the genome-encompassing capsid can be attached to a tail harbouring a host-recognition device called the baseplate (1) or they can exist without one, usually presenting spike protrusions on the surface of the capsid to allow for host recognition (2,3). While tailed phages have historically garnered the most attention, emerging metagenomic and environmental data are revealing a substantial array of tailless phages (4,5), with electron microscopy analysis of virion morphologies showcasing the potential dominance of these phages in diverse environments (6). The *Microviridae* family represents a major cohort within the realm of tailless phages. Notably, the *Microviridae* members are prominent constituents of aquatic habitats (7,8) and terrestrial animal gut viromes including those of humans (9). Variations in their abundance and diversity are linked to several disorders including inflammatory bowel disease, Crohn’s disease and cognitive dysfunction (10,11). The major capsid protein of *Microviridae* shares the single jelly-roll fold with capsid proteins from a wide range of viruses, including eukaryotic RNA and DNA viruses, some of which are major pathogens. The capsids shelter a circular ssDNA genome with sizes potentially ranging from 3.0 kb to 8.9 kb as suggested by metagenomic sequences (12,13). The genome usually encodes 6 to 10 proteins among which the major capsid protein VP1/F, DNA pilot protein VP2/H, internal scaffolding protein VP3/B, genome replication initiation protein VP4/A, and the ssDNA-binding protein VP8/J (14), stand out as most common.

Based on metagenomic data, the *Microviridae* family can be subdivided into over 20 groups, but only a handful of phages have been isolated and far fewer have undergone structural analysis (4). The two virus subfamilies with officially assigned taxonomies (15) are: the *Bullavirinae*, containing the model *Enterobacteria* phage phiX174; and *Gokushovirinae*, with best studied representatives including *Spiroplasma* virus SpV4 (16) and coliphage EC6098 (17,18). The key distinction between the subfamilies lies in their respective putative receptor-binding elements. *Bullavirinae* interacts with the host lipopolysaccharide (LPS) using a pentamer structure of the major spike protein G present at fivefold symmetry axes of its virion (2). In contrast, *Gokushovirinae* phages lack the major spike protein, opting instead for mushroom-shaped protrusions formed by the major capsid protein VP1/F at the threefold symmetry axes (16,18).

A molecular mechanism for initiating host cell infection has been previously proposed for the *Bullavirinae* phage phiX174 (2). According to this model, upon interaction with the LPS-core receptor, the pentameric structure of the major spike protein G dissociates from the virion. As a result, the hydrophobic loops of the major capsid protein become exposed and fuse with the LPS. Subsequently, the DNA pilot protein subunits located inside the virion rearrange, forming an oligomeric tube which is thought to create a channel for genome delivery into the host cytoplasm (19). However, this mechanism remains speculative as it was derived primarily from *in vitro* experiments. For *Gokushovirinae* and other subfamilies, the mechanisms of host recognition and genome delivery remain unknown (16,18).

Here, we report the structural analysis of the genome delivery process of a *Microviridae* phage, Ebor, along with its discovery and characterization. This phage, infecting a model aquatic Alphaproteobacterium *Rhodobacter capsulatus*, belongs to the microvirus taxon ‘Tainavirinae’ (20) and recognizes the host via five *Gokushovirinae*-like trimeric protrusions. Subtomogram averaging unveiled a mechanism whereby a capsid pentamer, nestled between trimeric protrusions interacting with the host cell, opens up and peels back from the centre. We referrer to this mechanism as the “stargate” because the capsid opening is reminiscent to the unique stargate vertex opening in giant viruses (21,22). These insights, integrated with previously published structural data on phiX174 (2,19), indicate that while different *Microviridae* subfamilies evolved to utilize distinct receptor-binding elements to recognize host cell, they all employ a conserved stargate mechanism of capsid opening to deliver genomic DNA during infection. The stargate mechanism of genome delivery, reported here, is potentially applicable across a broad spectrum of viruses that utilize single jelly-roll fold capsid proteins.

## Results

### Discovery of a model ‘Tainavirinae’ microvirus, Ebor

We decided to establish a tractable model system for *Microviridae* that would enable us to study the mechanism of genome delivery. The *Rhodobacter capsulatus* strain DE442, a mutant known for production of elevated amounts of a phage-like gene transfer agent (23), spontaneously produced phage particles that formed plaques on the parental strain *R. capsulatus* SB1003 (24). Pure phage DNA was isolated and sequenced using the Oxford Nanopore platform, revealing a 6611 bases-long single-stranded DNA genome. To our knowledge, this constitutes the largest isolated microvirus genome reported to date (**Figure 1a**), making it the most suitable system for structural analysis of ongoing host cell infection. The phage was named Ebor after the abbreviated Roman designation of the city of York, where this phage was isolated and described. The Ebor sequence was not identified in any of the complete genome sequences available in GenBank. However, sequence alignment with WGS genome entries of *Rhodobacter* strains, confirms that Ebor is transiently integrated as a prophage within the RuBisCO cbbII operon of *R. capsulatus* DE442, at the same site used by the dsDNA membrane-containing prophage Jorvik (3). Like in *Microviridae* and other ssDNA prophages, it is flanked by XerCD-binding *dif* motifs (14,25) (**Figure 1b**). The prophage was also detected in chromosome contigs of DE442-related strains Y262 and R121 (26) but is absent from the parental strain SB1003, and its precursor, the wild-type strain B10 (27). Therefore, we conclude that Ebor had to be acquired from a laboratory environment during the construction of gene transfer agent overproducers in the 1970s (28).

**Figure 1:**
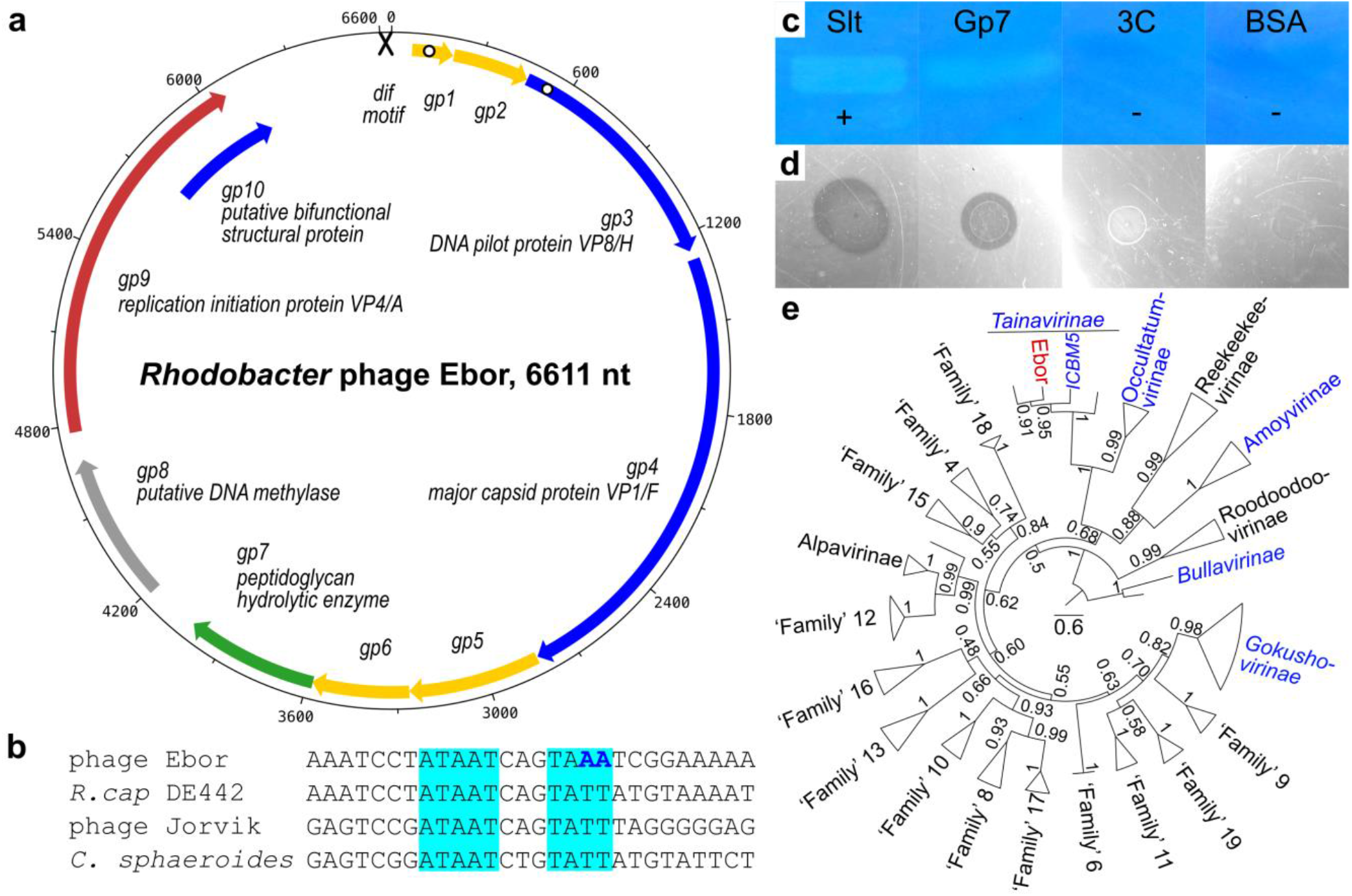
Overall characterization of the *Microviridae* phage Ebor. a) Genome map of phage Ebor. Genes encoding structural proteins (blue), lysin (green), replication initiation protein (red), methylase (grey), and unknown function (yellow) are depicted. The cross depicts the *dif* motif, white dots designate predicted transmembrane regions. b) Alignment of putative XerCD/dif motifs of phage Ebor, DE442 *Rhodobacter capsulatus* host bacterial strain, *R. capsulatus* phage Jorvik (3) and experimentally verified motif from related species *Cereibacter sphaeroides* (70). The canonical *dif* AT repeats are. highlighted in cyan, different nucleotides in the imperfect motif of Ebor are in bold blue. c-d) Zymogram (c) and water agar assay (d) of purified Gp7. Slt, a lysin from *Rhodobacter* phage Jorvik (3); 3C, 3C protease; BSA, bovine serum albumin; plus and minus signs denote positive and negative controls respectively. e) Phylogeny of conserved major capsid protein calculated using sequences from up to three representatives from each major lineage, denoted by triangles. Taxa lineages with isolated members are in blue, Ebor is highlighted in red. Numbers by nodes represent confidence values correspond to 100 transfer bootstrap estimates. The scale bar denotes amino acid substitutions/site.

We identified ten open-reading frames in the genome of Ebor (**Figure 1a, Table 1)**. These include the three most conserved genes of the *Microviridae* family: replication initiation protein VP4/A, DNA pilot protein VP2/H, and major capsid protein VP1/F (29). Mass spectrometry confirmed the presence of DNA pilot protein and major capsid protein in the sample of purified virions (**Supp Data S1**). The only other protein identified in significant amounts was the product of the *gp10*. This gene overlaps with the 3′-end of the gene encoding replication initiation factor VP4/A.

**Table 1:**
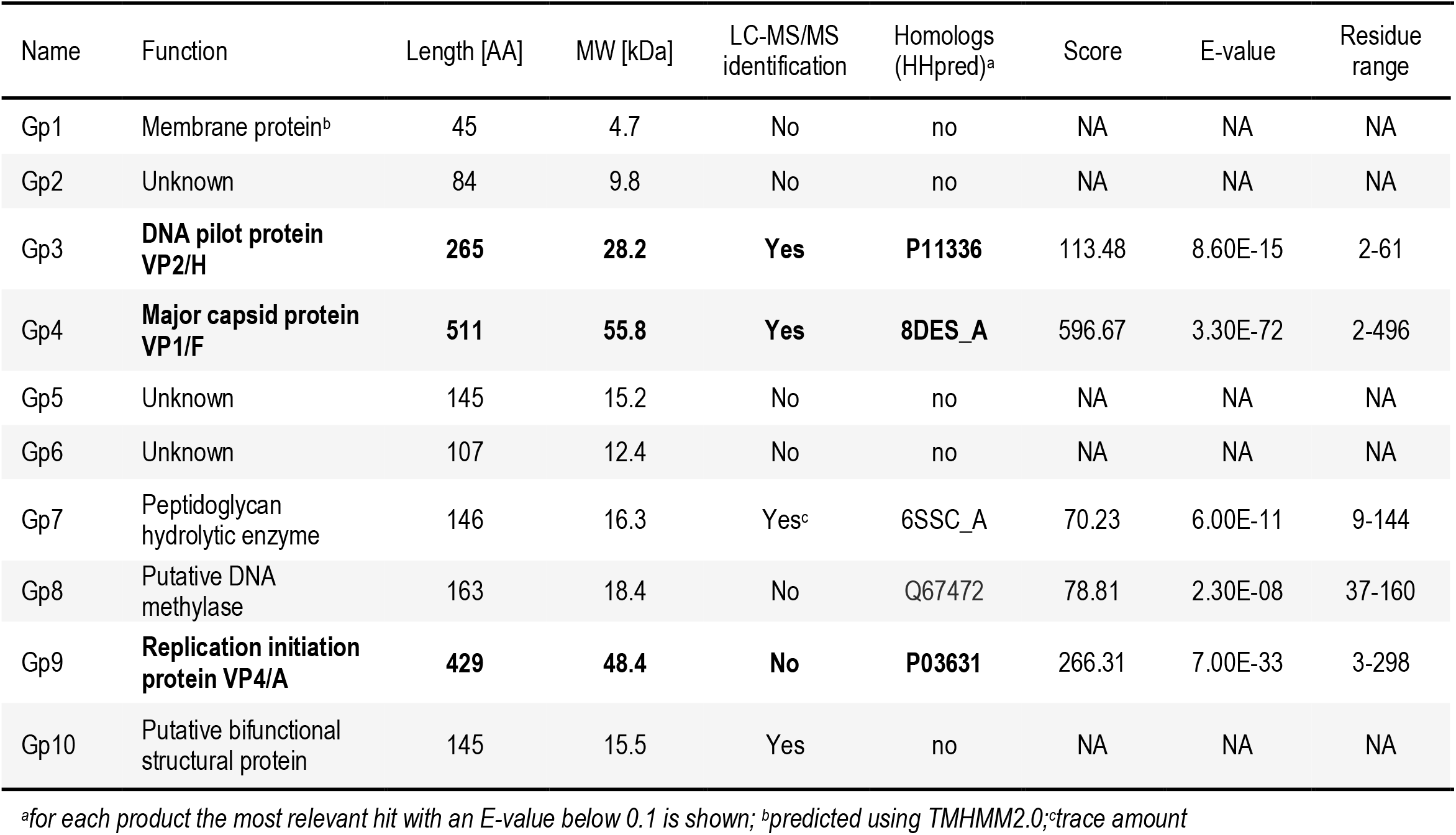
Gene products encoded by phage Ebor. The three signature products of the *Microviridae* family are highlighted in bold.

In addition to structural proteins, Ebor encodes Gp7 and Gp8. These proteins possess similarities to peptidoglycan hydrolytic enzymes and a probable DNA methylase, respectively, based on the primary sequence alignment and subsequent comparison of profile hidden Markov models by HHpred (30) (**Table 1**). Purified Gp7 degrades crude *Rhodobacter* peptidoglycan in zymogram and water agar assays (**Figure 1c-d, Supp Figure S1**), and to our knowledge, Ebor represents the first experimentally verified ssDNA phage possessing a peptidoglycan hydrolytic enzyme. According to analysis of nanopore sequencing data using the modkit tool, 11.53% of adenine calls belonged to 6mA, 1.58% of cytosine calls to 5mC and 0.97% to 5hmC in any contexts. This proves Ebor genomic DNA is methylated; however, we cannot assess if the methylation is performed by Gp8 or host methylases.

Taxonomic analysis with MOP-UP, a protein sharing network algorithm using an up-to-date database of *Microviridae* genomes (4) (**Supp Data S2**), placed Ebor within the recently proposed candidate subfamily ‘Tainavirinae’ (20) and in a genus-level grouping with other *Rhodobacteraceae* prophages, which we confirmed by phylogenetic tree reconstruction (**Figure 1e**). The genome composition of Ebor is typical of the ‘Tainavirinae’ subfamily except for the putative methylase gene, which is only sporadically present in *Microviridae*. We note significant differences between ‘Tainavirinae’ and isolates from other *Microviridae*. The only shared proteins with recognizable homology are the major capsid protein VP1/F, replication initiation protein VP4/A and DNA pilot protein VP2/H. However, the amino acid identity for these shared proteins is below 30 % (**Supp Figure S2**).

### Extended protrusions at 3-fold symmetry axes of the capsid unfold during genome ejection

We isolated two Ebor variants exhibiting different stability during cryo-EM sample preparation. Nanopore sequencing showed a single nucleotide polymorphism leading to S120R amino acid replacement in the major capsid protein. Thus, we designated these variants S120 and R120. Variant R120 virions were purified using CsCl gradient, vitrified and imaged using cryo-EM, yielding stable native particles. Single particle reconstruction produced an icosahedrally averaged map of Ebor at 3.2 Å resolution (**Figure 2a, Supp Table S1**). The capsid shell has an external diameter of 28 nm and an internal diameter of 25.1 nm. The internal diameter is 3.2 to 4.6 nm larger compared to other determined capsid structures of *Microviridae* members, allowing Ebor to accommodate larger genome (**Supp Figure S3**). The capsid exhibits the triangulation number of T=1, being made of 60 copies of the major capsid protein (**Figure 2b**). The core of the major capsid protein contains an 8-stranded antiparallel β-barrel (single jelly-roll fold), the fold shared among all *Microviridae* (**Figure 2b-c; Supp Figure S3**). A striking characteristic of the capsid is the presence of elongated protrusions situated at its three-fold symmetry axes, reaching out 9 nm from its surface. These extensions are formed by loops composed of residues 184-254 from three neighbouring subunits of the major capsid protein (**Figure 2a-b**). This is the same loop of the major capsid protein that forms trimeric protrusions in *Gokushovirinae* members (16,18). As the density of the protrusions was resolved to a lower resolution, a model of this loop generated using AlphaFold2 was fitted into the experimental map (**Figure 2d, Supp Data S3**). The model showed the tip of the protrusion possesses a positive electrostatic potential, contrary to the negative potential of the capsid’s penton exterior (**Supp Figure S3**).

**Figure 2:**
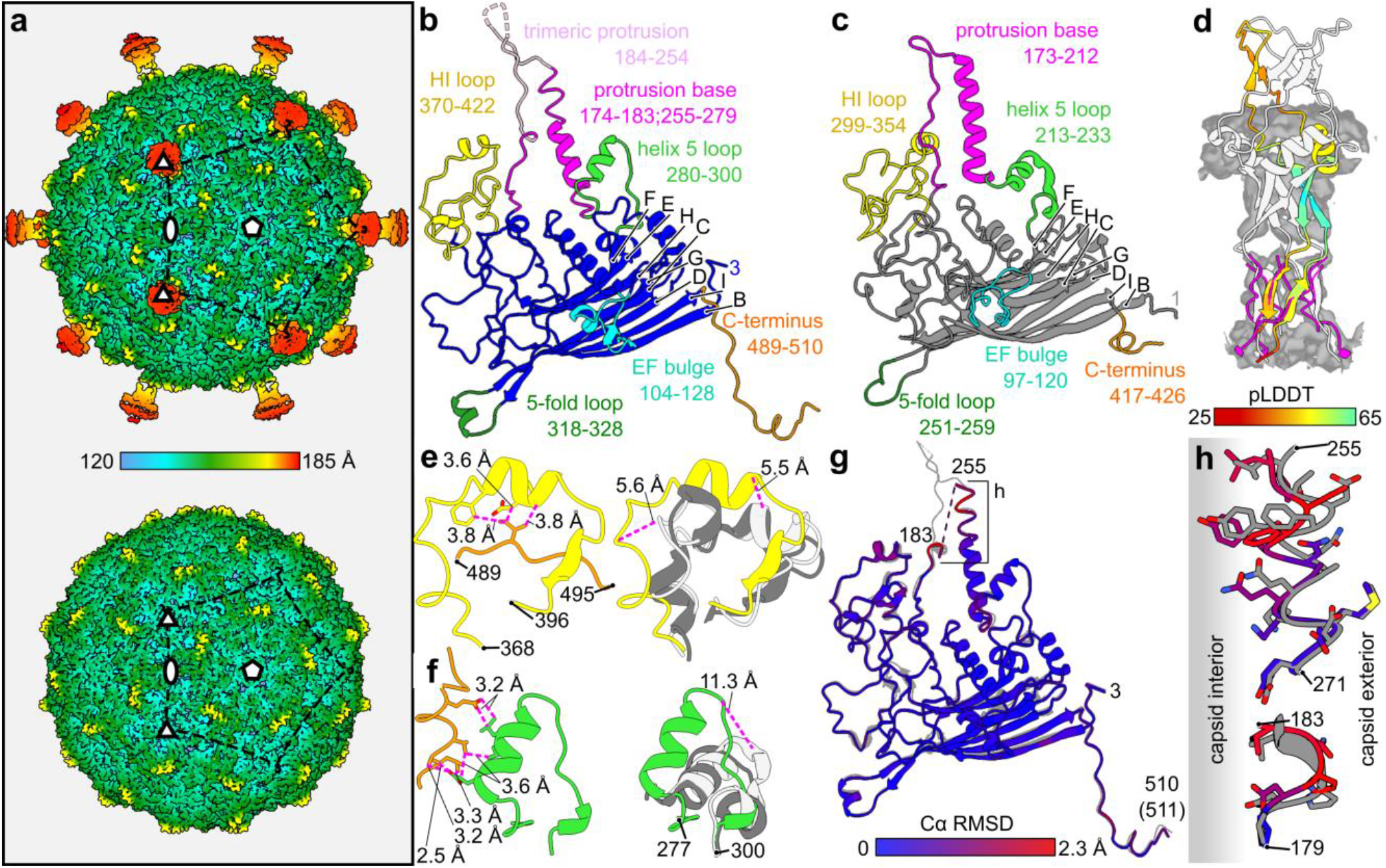
Structure of the Ebor virion determined by single particle analysis. a) Icosahedrally averaged map of the native capsid decorated with trimeric protrusions (top) and the empty capsid with unfolded protrusions (bottom). The symmetry axes and position of one pentamer are highlighted. The colour bar indicates the distance from the centre of the capsid. b-c) Ribbon diagrams of the major capsid protein of Ebor (b) and phiX174, PDB code 2BPA_B (c). β-strands are designated according to convention (31). Regions of interest are highlighted in colour, annotated with a description and delimiting residue numbers are shown. d) AlphaFold2 model of the protrusion (residues 184-254) fitted in the corresponding density (grey). One subunit is coloured according to the prediction confidence measure pLDDT. The built regions from b) are superimposed and shown in magenta. e-f) Interaction of HI loop (e) and helix 5 loop (f) with the C terminus (left) causes their misplacement from the structures of phiX174, PDB code 2BPA_B (light grey) and EC6098, PDB code 8DES_A (grey) as shown by their superimposition (right). The distances between the interacting residues of Ebor and corresponding Cα of phiX174 and Ebor models are shown. g) Superimposition of models built into the native (grey) and empty map (colour). The colour-coding is based on RMSD between corresponding Cα. h) Close up view of the protrusion base region where the largest deviations between the two models is present, shifting the residues to the capsid interior.

The C-terminus of the major capsid protein is extended compared to other known structures of *Microviridae* members. Positioned beneath the loop connecting β-strands H and I as specified in phiX174 (31) (referred within the rest of this paper as the HI loop) of the adjacent subunit, it resides above the base of the protrusion, interacting with the helix 5 loop previously reported for phiX174 to reversibly bind the host surface (32). These interactions with the C-terminus results in a distinct conformation of the HI loop and helix 5 loop, compared to proteins of phiX174 and EC6098 **(Figure 2e-f)**. Consequently, a single protein unit adopts a less compact structure, facilitating increase in the curvature of the capsid.

When a cryo-EM sample of the variant S120 was prepared using the same protocol described above, some particles on the grid were observed to lose their genomes (**Supp Figure S4**). Further analysis of the dataset showed that ∼10% of particles were missing one pentameric capsomer (**Supp Figure S3**). Subsequently, using a different purification method based on a sucrose gradient, we obtained a virus titre comparable to that obtained with CsCl gradient, as estimated by plaque assay. However, examination on the cryo-EM grid showed that all the particles had ejected the genome (**Supp Figure S4**). The ejection likely occurred during the blotting procedure due to the reduced stability of the S120 variant. The icosahedrally averaged reconstruction of the empty particles led to a 3.3 Å resolution map, which notably lacked the trimeric protrusions (**Figure 2a)**. Otherwise, only subtle differences were observed between the major capsid protein models, built into the maps of native R120 and empty S120 particles. The respective models exhibited an average RMSD_Cα_ of 0.55 Å with the maximal RMSD_Cα_ of 2.3 Å occurring at the base of the protrusion (**Figure 2g-h**). Here, the residues move towards the capsid interior as they are not repelled by the pressurized genome, thereby inducing the unfolding of the protrusion loop trimer.

### Ebor recognizes the host LPS as its receptor

We incubated susceptible host strains B10 and SB1003 (**Figure 3a-c**) with the stock of Ebor to assess the identity of the host receptor. In the case of B10, where Ebor produces larger plaques, a large number of phage particles bound to the cell membrane (**Figure 3d**). In the case of SB1003, where Ebor forms small plaques, almost no particles bound to the cells (**Figure 3e**). The addition of 5 mM Ca^2+^ increased the size of Ebor plaques on the SB1003 strain. When 5 mM Ca^2+^ was added to samples prior to vitrification, it resulted in a greater number of attached particles; however, all of these particles appeared to bind to the capsule rather than the cell membrane (**Supp Figure S4**). To test if capsular polysaccharide is required for infection, we subsequently infected SB1003Δ*gtaI*, a quorum sensing system mutant incapable of forming the capsule (33). The knockout strain exhibited larger plaque morphology than SB1003 (**Figure 3c**), with phage particles attached to the outer membrane, albeit in significantly lower numbers compared to B10 (**Figure 3f**). This indicates that the capsule is not essential for infection.

**Figure 3:**
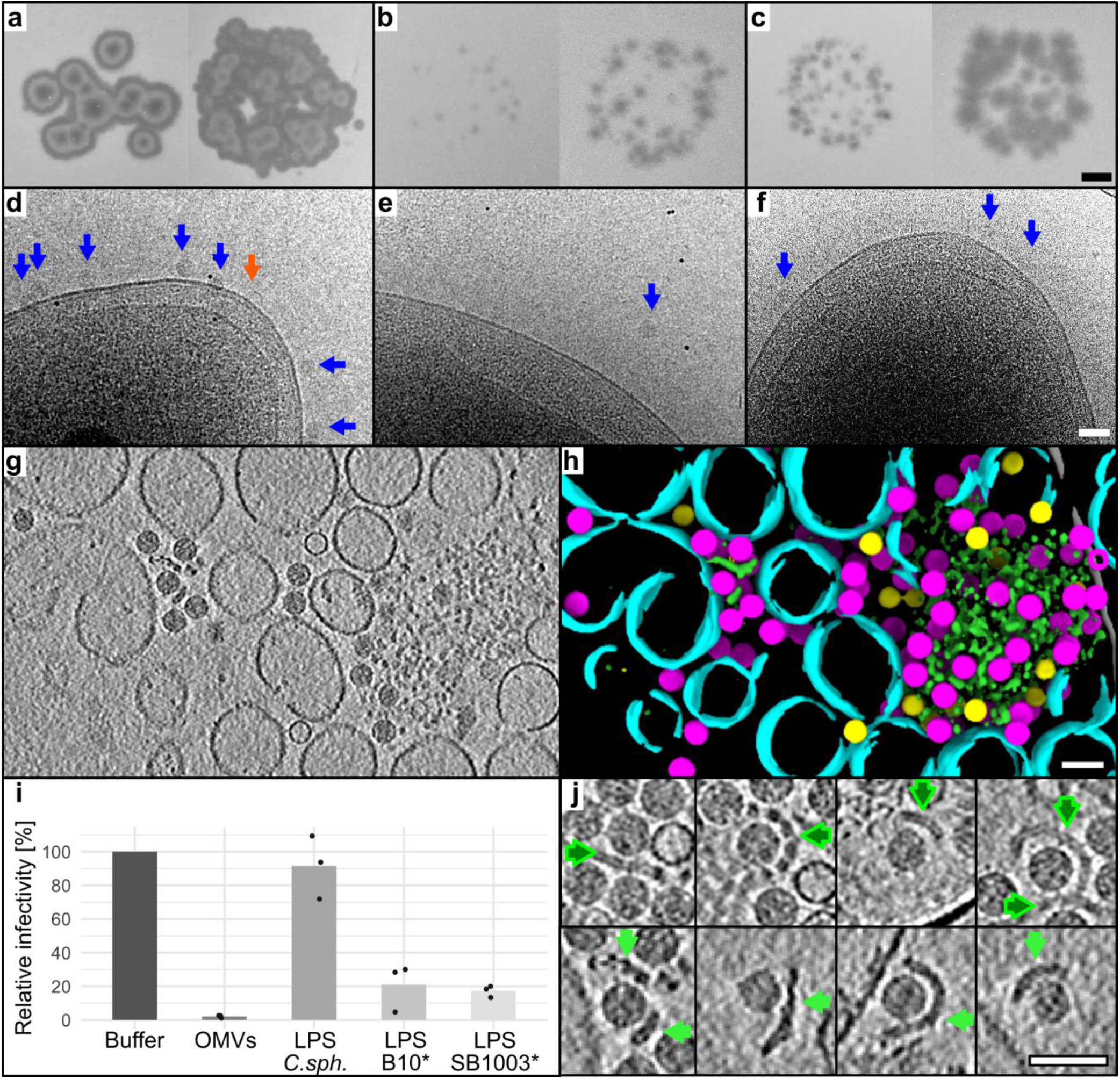
Characterization of the Ebor host receptor. a-c) The phage Ebor variant S120 (left panels) and R120 (right panels) were spotted onto soft agar plates containing the host strain B10 (a), SB1003 (b), and SB1003Δ*gtaI* (c). The scale bar is 2 mm. d-f) The stock of variant R120 purified by ion exchange chromatography was imaged by cryo-EM after 5 min of incubation with the host strain B10 (d), SB1003 (e), and SB1003Δ*gtaI* (f). The native and empty virions are depicted by blue and orange arrows, respectively. The scale bar is 50 nm. Images were collected at 200 kV using a Glacios TEM equipped with Falcon 4 camera, with the total exposure dose ∼3 e^−^/Å^2^. g-h) Central Z slice of a tomogram of Ebor virions mixed with purified OMVs (g) and its segmentation (h). Native capsids (magenta), empty capsids (yellow), single layered vesicles (cyan), and bilayered polymer (green) are highlighted. The scale bar is 50 nm. i) Phage infectivity after incubation with different putative receptors. LPS, lipopolysaccharide; OMV, outer membrane vesicle; C.sph, *C. sphaeroides*; *, Δ*gtaI* strains. Individual measurements of biological replicates are plotted as points, their averages are plotted as bars. The raw values are shown in **Supp Table S2**. j) Cryo-EM of four examples of phage particles attached to LPS bilayers isolated from B10Δ*gtaI* (dark green arrows) and bilayered polymers present in OMV sample (light green arrows). The scale bar is 50 nm.

To further investigate the identity of the Ebor receptor, we mixed the virus with purified outer membrane vesicles (OMVs) formed from disrupted B10 cells. Cryo-ET showed that the OMV sample consisted of single-layered vesicles 50 to 150 nm in diameter, larger single-layer membrane patches and bilayered polymers that tended to aggregate and attracted most of the phage particles (**Figure 3g-h; Supp Movie S1**). As LPS forms bilayers, can aggregate and is a demonstrated receptor for phiX174 (2,34), we isolated LPS from *R. capsulatus* and tested its inhibition effect on Ebor. The *gtaI* knockout strains were selected to ease the purification of the LPS from capsular polysaccharide. After ten minutes, Ebor titre dropped by 97.9 % when incubated with OMVs and approximately 80 % when incubated with LPS (**Figure 3i, Supp Table S2**). Cryo-ET showed that the LPS sample comprised bilayers that attracted phage particles, closely resembling the shape observed in the bilayered polymer present in the OMV sample (**Figure 3j)**. LPS extracted from B10Δ*gtaI* and SB1003Δ*gtaI* exhibited similar banding patterns on polyacrylamide gels (**Supp Figure S1**) and inhibited the phage to a similar extent (**Figure 3i**). These similarities do not explain the different attachment abilities of Ebor on these strains. Considering the relatively weakly inhibitory effect of purified LPS, it is possible that other cell surface polymers are involved in Ebor’s host recognition.

### Ebor binds to host via five protrusions, inducing capsid penton opening

We reconstructed tomograms of Ebor attached to native B10 cells (**Figure 4a-b, Supp Movie S2**). Subtomogram averaging of the 1593 attached particles revealed that the phage binds to the outer membrane using five protrusions, causing the penton nestled between them to approach the host surface (**Supp Figure S5**). This attachment pattern differs from that observed for particles attached to OMVs, where only two protrusions were involved (**Supp Figure S6**), likely due to differences in membrane curvature. By performing focused classification around the interacting region, we identified three different states of the particles, with the interacting penton moving closer and ultimately fusing with the membrane (**Figure 4c**). To enhance the resolution of these intermediates and validate their existence through an alternative method, we collected a single particle dataset on the same sample, targeting acquisition areas (35) on cells. This yielded 23095 attached particles with correctly estimated membrane orientation (**Supp Figure S7**). Within this subset, we identified three classes corresponding to those obtained by subtomogram averaging (**Supp Figure S8**).

**Figure 4:**
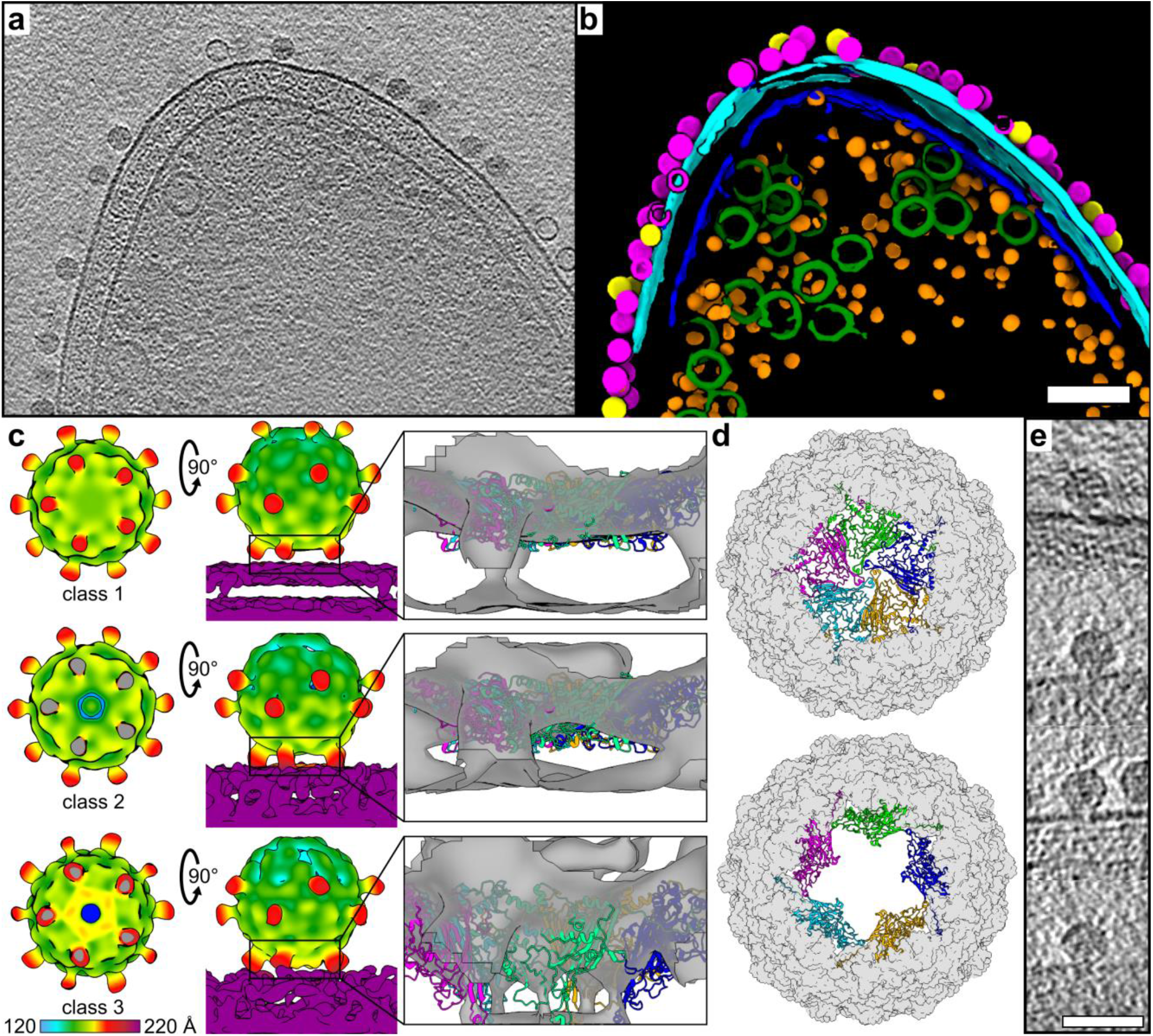
Cryo-ET analysis of Ebor interaction with the host cell. a-b) Central Z slice of a tomogram of an *R. capsulatus* strain B10 cell with attached Ebor virions (a) and segmentation of 127 nm-thick slab of the respective tomogram (b). Native (magenta) and empty (yellow) capsids, outer membrane (cyan), inner membrane (blue), intracytoplasmic membrane (green) and ribosomes (orange) are highlighted. The scale bar is 100 nm. c) C5 symmetry maps of the attached particles reconstructed by subtomogram averaging, classified into class 1 (top), 2 (middle) and 3 (bottom). Top and side views are shown along the interacting pentamer of the major capsid protein fitted into the unfiltered map of each class. In the top view, the membrane is cropped out with grey indicating virion-membrane connection sites where virion density has also been cropped. In the side view, the upper trimeric protrusions are not always visible in the set density threshold due to ice thickness gradient but actually present in all maps. The colour bar indicates the distance from the centre of the capsid. d) Models of capsids generated by fitting capsid protein subunits into the maps of class 1 (top) and class 3 (bottom). Capsids are viewed towards the penton that opens up, with the central penton shown as ribbons (subunits in different colours) and the rest of the capsid shown as a molecular surface (grey). e) examples of Z slices of particles from polished high-defocus tomograms that shows penton opening towards the cell membrane. The scale bar represents 50 nm.

The first class represents Ebor attached to cells where no conformational change of the virion core compared to the particle in solution occurs (**Figure 4c, Supp Figure S8**). The density of protrusions is only weakly connected with the membrane. In the second class, the trimeric protrusions are in closer contact with the membrane, with a pore forming along the 5-fold axis of the penton nestled between the protrusions (**Figure 4c, Supp Figure S8**). A comparable classification of weak and strong connection of protrusions to the membrane density was observed for particles interacting with OMVs (**Supp Figure S6**). The third class (class 3) showed the penton opening further, fusing with the membrane, and generating a pore extending from the capsid (**Figure 4c, Supp Figure S5**).

Maps of class 3 generated by the two methods differed slightly from each other. The map generated by single particle analysis had an extra density connecting the virion with the membrane, corresponding to the position of the EF bulge, with the central part of the interacting penton density being distorted (**Supp Figure S8**). This map strikingly resembles that of phage phiX174 interacting with LPS (2). It is important to note that the EF bulge of Ebor lacks a hydrophobic region conserved within *Bullavirinae* members, likely as a hydrophobic loop exposed at the virion surface would be detrimental to its stability. The only exposed hydrophobic residue of the loop is L124 (**Supp Figure S8**). In comparison, the map generated from subtomogram averaging showed a more pronounced bulging of the interacting penton into the membrane, suggesting adjustments in the position of entire subunits of the pentamer. Fitting the capsid protein into this density results in a virion model with a stargate opening of the interacting penton. In this model, the major capsid protein subunits forming the pentamer begin to separate from each other, gradually peeling away from the centre of the pentamer (**Figure 4d, Supp Movie S3-4**). This motion is analogous to the observed capsid opening in giant viruses where it occurs at the unique 5-fold vertex (21,22). We note that a pronounced bulging, consistent with the stargate mechanism of capsid opening, was also visible in single particles upon inspection of the polished tomograms (**Figure 4e**).

## Discussion

The identification of Ebor, the largest microvirus characterised to date, enabled accurate structural characterisation of host infection by cryo-ET. Ebor represents a member of a distinct subfamily within *Microviridae*, exhibiting a significant evolutionary divergence from the previously characterized *Bullavirinae* phage phiX174. This divergence is exemplified by the considerably larger genome, totalling 6.6 kb compared to phiX174’s 5.4 kb. To accommodate it, Ebor utilises an expanded capsid with ∼83 % larger internal volume, built with the help of an extended C-terminus of the major capsid protein and several loops adopting a stretched conformation. The larger genome accommodates two genes that are not universally conserved within *Microviridae*: *gp7*, bearing resemblance to genes encoding peptidoglycan hydrolytic enzymes found in tailed phages, and *gp8* that encodes putative DNA methylase and likely contributes to phage resistance against host defence systems. We showed that Ebor’s genome is predominantly methylated at adenines (6mA); however, we note the real level may differ as the methylation detection methodology hasn’t been trained on bacteriophage genomes. In addition, other types of modifications may be present that are not currently detectable by the Nanopore platform (36). Another notable feature of Ebor is the presence of the virion-associated protein Gp10 that is conserved within the ‘Tainavirinae’ subfamily. Its genome location and size is similar to the *Bullavirinae* scaffolding protein B (**Supp Figure S2**). However, Gp10 is the only protein identified as a virion component that is highly positively charged with the theoretical pI of 11.3. It thus may have an analogous function to the conserved negative charge-mitigating ssDNA-binding protein VP8/J, which replaces the scaffolding protein as the ssDNA genome is loaded into the capsid of both *Bullavirinae* (37) and *Gokushovirinae* (17).

Based on structural findings, we postulate the following mechanism for phage Ebor genome delivery (**Figure 5**). First, Ebor recognizes cell surface polysaccharides, including capsular polysaccharide and LPS; however, unlike the *R. capsulatus* gene transfer agent and related bacteriophages, the capsular polysaccharide is not an essential receptor (23). Second, Ebor’s irreversible attachment is mediated via five trimeric protrusions of the major capsid protein that bring the phage closer to the outer membrane. This interaction instigates formation of a pore along the 5-fold symmetry axis of the interacting penton. Finally, the stargate opens up, facilitating genome delivery. Given the rigidity of the β barrel forming the core of the major capsid protein subunit, it is reasonable to assume that the stargate opening occurs through the disruption of subunit-subunit interactions and their separation within the pentamer, while each subunit maintains interactions with the rest of the capsid. The energy required for this opening could potentially be generated by favourable interactions of the trimeric protrusions with the host’s membrane and entropic gains associated with the release of coordinated water molecules.

**Figure 5:**
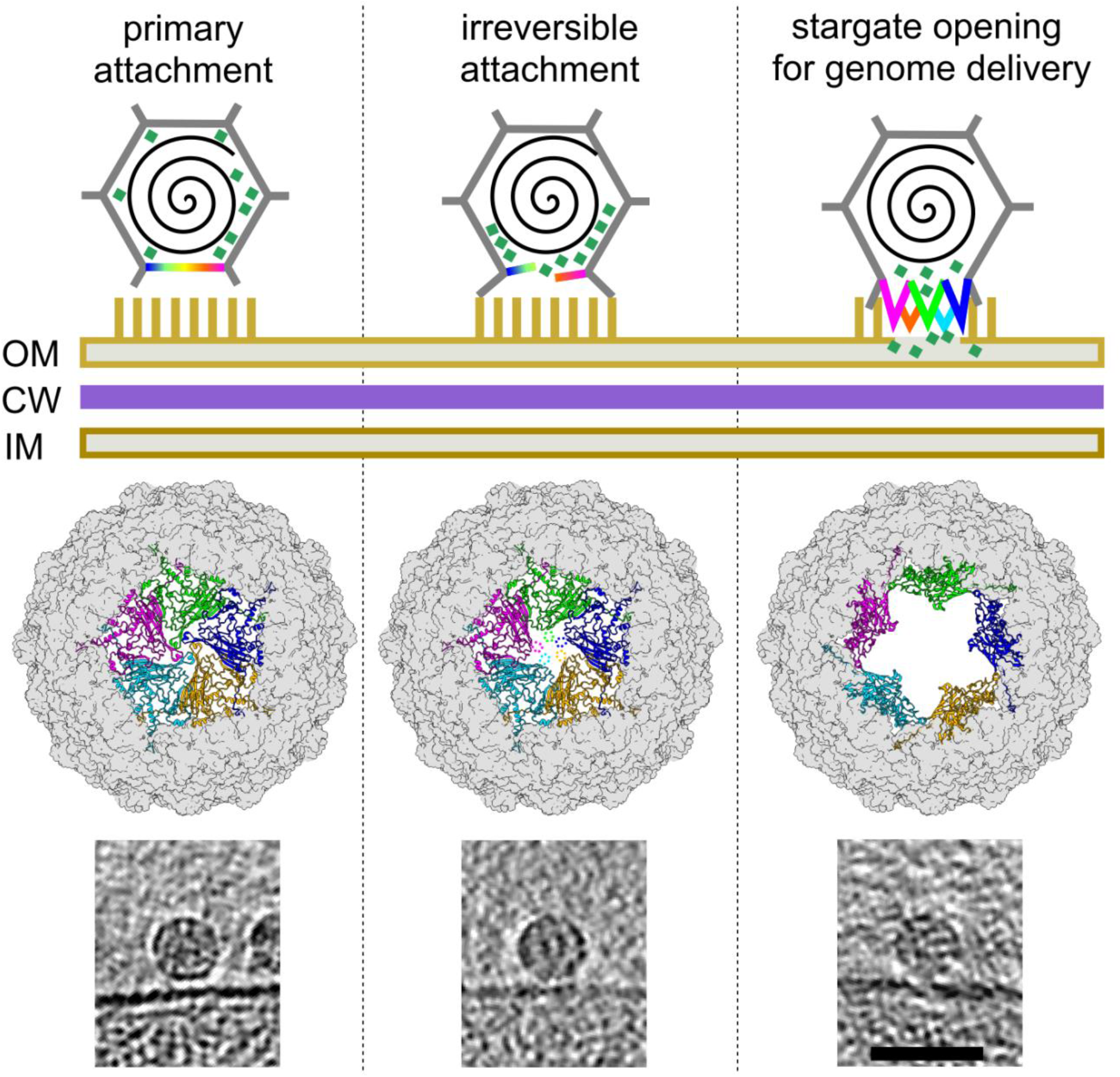
Proposed genome delivery mechanism of phage Ebor. The three specific states, labelled at the top, correspond to classes 1-3 shown in **Figure 4c**. Top row: A graphic illustration depicting three specific states of infection initiation, showing the virion (grey), interacting penton (rainbow), genome (black spiral), internal DNA pilot proteins (aquamarine squares), and lipopolysaccharides (yellow sticks). OM, outer membrane; CW, cell wall; IM, inner membrane. Middle row: capsid viewed towards the penton that opens up, with the central penton shown in ribbons and the rest of the capsid shown as a grey molecular surface. Each major capsid protein subunit is depicted in different colour. Bottom row: examples of Z slices of particles from polished high-defocus tomograms corresponding to the three individual steps. The scale bar is 50 nm.

Notably, both maps of the class 3 genome-releasing state (class 3), derived from subtomogram averaging (**Figure 4c**) and single particle analysis (**Supp Figure S8**), showed that the virion’s penton fuses with the membrane. The map derived from subtomogram averaging aligns best with the stargate model, containing density for complete major capsid protein subunits. In the map derived from the single particle analysis a vast portion of the penton density, including the highly stable core β barrel, appears weak and distorted. The inferior resolution is likely caused by the single particle acquisition of viruses attached to cells, being more prone to bias from preferential orientations. This bias arises because only particles perpendicular to the incident electron beam could be analyzed due to cell thickness. Similar bias was reported for the dataset of phiX174 attached to LPS (2) with the derived map of the genome delivery intermediate also exhibiting a loss of the signal around the central penton, preventing precise localization of respective major capsid protein subunits.

In phiX174, the internal DNA pilot proteins multimerize to form a DNA delivery tunnel (19). It was also shown that the genome-releasing intermediate contains a cavity with weaker density inside the capsid, hypothesized to be caused by the rearrangement of pilot proteins (2). We observed a weaker internal density right above the interacting pentamer in classes 2 and 3, suggesting ongoing rearrangement (**Supp Figure S8**), albeit less prominent than in phiX174 where the void cavity extends further and reaches the centre of the capsid. Furthermore, classes 2 and 3 show an increased density of the membrane at the site of penton fusion, which bulks towards the interacting penton in class 3 (**Figure 4c, Supp Figure S8**). This could be attributed to the insertion of the hydrophobic N-termini of DNA pilot proteins. Subtomogram averaging and inspection of polished tomograms did not identify pronounced periplasmic tunnels, as seen for phiX174 infecting *E. coli* minicells (19). Therefore, the mechanism by which the Ebor genome is translocated through the periplasm remains be uncovered.

We conclude that genome delivery within the *Microviridae* family occurs via a general stargate mechanism of capsid opening along the 5-fold symmetry axis, with LPS serving as a broadly recognized receptor. The stargate opening of the capsid has been previously proposed for giant dsDNA viruses, where it occurs on a different molecular scale, but serves the same purpose of delivering the viral genome (21,22). Utilization of the same mechanism by the *Microviridae* family underscores the universal applicability of the stargate opening across different virus realms. It may potentially also apply to other viruses composed out of single jelly-roll capsid proteins such as *Picornaviridae* (38), providing an alternative explanation on how picornaviruses lose the pentameric capsomer observed during *in vitro*-mimicked infection conditions (39).

## Materials and methods

### Bacteria, media and phages used in this study

The bacterial strains used in this study are summarized in **Supp Table S3**. *R. capsulatus* strains were grown either in minimal medium RCV (40) or rich medium YPS (41), aerobically at 30°C. When working with the phage Ebor and strain SB1003, an additional 0.002 M CaCl_2_ was added to the liquid and soft agar media unless otherwise stated.

### Isolation of the phage Ebor and generation of phage variants

*R. capsulatus* DE442 was incubated in RCV medium at 30°C/200 rpm for 30 hours, reaching a late log phase. The cells were pelleted at 4000 × g for 6 min and 100 µl of the supernatant were added to 100 µl of stationary phase *R. capsulatus* SB1003. The mixture was overlayed onto YPS plates and incubated overnight at 30°C. A single plaque was picked using a sterile tip, resuspended in 100 µl of YPS medium and further propagated on a plate using the same propagation strain. After the second round of propagation, a spontaneous clear plaque appeared and was propagated on plates using strains SB1003 and B10 respectively. After several rounds of propagation on strain B10, the phage stopped producing clear plaques on strain SB1003. This phage was designated as variant S120, while the phage propagated continuously on SB1003 was designated as variant R120.

### Production of Ebor in a high titre

Phage Ebor was washed from three confluently lysed plates using 5 ml of YPS, and this phage-containing medium was added to 25 ml of the propagation strain (SB1003 for variant R120, B10 for variant S120) grown to OD_600nm_ = 0.15. The mixture was incubated for 7 hours at 26°C, 140 rpm. The supernatant was then added to 250 ml of the propagation strain grown to OD_600nm_ = 0.15 and incubated in the same manner. After this incubation, the titre of the lysate was in the range of 10^9-10^ PFU/ml. The lysate could be aliquoted to 0.5 ml and stored at -80°C for a year without an apparent loss of the titre.

### OMV extraction

The procedure was modified from Cian et al. (3). *R. capsulatus* was grown in 1 L of 1:1(V:V) YPS:RCV medium (42) to stationary phase and pelleted by centrifugation at 4°C, 6900 × g for 15 min. The pellet was resuspended in 12.5 ml of 0.5 M sucrose in 10 mM Tris-HCl, pH = 7.5, supplemented with 114 *µ*g/ml lysozyme and incubated on ice while stirring for 2 min; 12.5 ml of 1.5 mM EDTA were added and stirred for 7 min. The mixture was centrifuged at 4°C, 11400 × g for 10 min and the pellet was resuspended in 25 ml of 0.2 M sucrose, 10 mM Tris-HCl, pH = 7.5, 2.2 mM MgCl_2_, 25 ng/ul DNase I with the addition of 1 ul of EDTA-free protease-inhibitor cocktail (MilliporeSigma). The mixture was homogenized with a Dounce homogenizer and lysed in French press (Aminco) at 4°C, 4 cycles at 20,000 psi. The lysate was centrifuged at 4°C, 6700 × g for 10 min, and the supernatant was further centrifuged at 4 °C, 42,400 rpm for 90 minutes (Beckman 70Ti). The pellet was resuspended in 1 ml of 10% sucrose solution using a Dounce homogenizer and put on a step gradient of 2 ml of 73% sucrose, 4 ml of 53% sucrose and 6 ml of 20% sucrose. The gradient was centrifuged at 4 °C, 34000 rpm for 45 h (Beckman SW-41) and the lowest band collected, which was loaded on a second identical sucrose gradient for maximum removal of intracytoplasmic membrane (ICM) traces from the OMV fraction. The lowest band was collected once again and diluted 1:1 in 10 mM Tris, pH = 7.5 and pelleted at 4 °C, 70000 rpm for 60 min (Beckman TLA-100.3). The pellet was resuspended in 500 µl of 10 mM Tris, pH=7.5 and kept on ice.

### LPS extraction

The procedure was based on Lam et al. (4). *R. capsulatus* grown in 1 L of 1:1 (v/v) YPS:RCV medium to stationary phase was harvested by centrifugation at 4°C, 6900 × g for 15 min. Pellets of approximately 1.5 g wet weight were resuspended in 10 ml of 100 mM NaCl and heated to 68°C. Ten ml of phenol prewarmed to 68°C were added and the mixture was vortexed vigorously every 15 min for 1 h at 68°C. The mixture was cooled on ice for 10 min and centrifuged at 4°C, 12000 × g for 15 min. The upper phase was transferred to a clean container. Ten ml of 100 mM NaCl were added to the lower phase and the 68°C incubation and centrifugation steps were repeated. The pooled upper phases were dialysed (in 3.5 kD cutoff tubing) against 3.5 L dH_2_O overnight and a second 3.5 L dH_2_O for 2 h. The sample was supplemented with MgCl_2_ (5 mM), DNase I (20 *µ*g/ml) and RNase A (20 *µ*g/ml), and incubated at 37°C for 1 h. Proteinase K was added (30 *µ*g/ml) followed by incubation at 50°C for 2 h. The sample was lyophilized and resuspended in 4 ml of 20 mM NaOAc, pH 7.5, and centrifuged at 4°C, 200000 × g overnight. The pellet was resuspended in 0.5 ml of 10 mM Tris-HCl (pH 7.5), 0.1 mM MgCl_2_.

### Purification of Ebor using ultracentrifugation

From 500 ml of phage lysate containing 10^9-10^ PFU/ml, cells and cell debris were removed by centrifugation at 8000 × g for 30 min. NaCl and PEG8000 were added to the lysate to a final concentration of 0.5 M and 8% w/v respectively, dissolving at 4°C for 1 hour. The sample was centrifuged at 4°C, 14 000 × g for 25 min, and the pellet was resuspended in 15 ml of E-buffer (20 mM Tris, pH 7.75; 100 mM NaCl; 5 mM CaCl_2_). For the caesium chloride ultracentrifugation, the sample after PEG precipitation was first spun at 10°C, 35000 rpm for 3.5 h (Beckman Ty70.1Ti). The pellet was resuspended in 0.7 ml of E-buffer overnight and loaded onto handmade 1.3-1.4-1.45-1.5-1.7 g/ml caesium chloride density gradients. The sample was centrifuged at 12°C, 21000 rpm for 5 h (Beckman SW41Ti). The band of the expected density was pooled using a syringe and the buffer was exchanged to the E-buffer using 100kDa Vivaspin^TM^ columns (Cytiva). This method resulted in the production of purified variant R120 showing a titre in the range of 10^10-11^ PFU/ml that showed stable virions after vitrification; and genome-releasing virions of variant S120 showing a titre 3 × 10^10^ PFU/ml (**Supp Figure S4**).

For the sucrose gradient centrifugation, the sample after PEG precipitation was loaded onto 15-30 % sucrose gradients prepared using Gradient Master^TM^ (Biocomp) and spun at 15°C, 22000 rpm for 1.5 h using Beckman SW28 rotor. The fractions containing the phage were pooled and spun at 10°C, 35000rpms for 3.5 h (Beckman Ty70.1Ti). The pellet was resuspended in 0.7 ml E-buffer overnight and loaded into 20-40% sucrose gradients prepared using Gradient Master^TM^. The sample was centrifuged overnight at 10°C, 22 000 rpm (Beckman SW41Ti) with the phage sedimented at the bottom of the tube. The fraction was taken by cannula syringe and the buffer was exchanged to the E-buffer using 100kDa Vivaspin^TM^ columns. This method resulted in the production of purified variant S120 reaching the titre of 5 × 10^10^ PFU/ml that upon vitrification contained only empty particles (**Supp Figure S4**).

### Purification of Ebor using ion exchange chromatography

From 500 ml of phage lysate, cells and cell debris were removed by centrifugation at 8000 × g for 30 min. The supernatant was consecutively filtered through 0.8 and 0.4 μm filters. The filtered supernatant was mixed with the loading buffer (20 mM Tris, pH 7.75; 10 mM NaCl; 5 mM CaCl_2_) in the ratio 1:2 v/v and applied to a pre-equilibrated CIM-multus QA 8 ml column (BIAseparations). Particles of virus bound to the column were washed by applying 12 column volumes of the loading buffer followed by a linear gradient of elution buffer (20 mM Tris, pH 7.75; 1.8 M NaCl; 5 mM CaCl_2_) until the conductivity of the mixture reached 32 mS (∼14% v/v). The elution of virus particles was induced by an increasing concentration of the elution buffer until the conductivity reached 40 mS (∼16% v/v). The fractions containing the virus were identified using titering and pooled. The concentration of NaCl in the pooled sample was increased to 0.5 M and 8 % w/v of PEG8000 was added, dissolving at 4°C for 1 hour. The sample was centrifuged at 4°C, 14 000 × g for 25 min, and the pellet was resuspended in the E-buffer. The sample was then aliquoted, frozen in liquid nitrogen and kept at -80°C. The sample showed a titre of 3 × 10^11^ PFU/ml and when imaged a mixture of native and empty particles in a similar ratio (**Supp Figure S4**).

### Ebor inhibition assay

The concentration of LPS isolated from B10ΔgtaI and SB1003ΔgtaI, and purchased LPS from *Cereibacter sphaeroides* (InvivoGen) was estimated based on the intensity of the LPS-corresponding band from the individual samples that were ran on a polyacrylamide gel stained using silver stain kit (Thermo Fisher Scientific) and alcaine blue (Millipore Sigma) (**Supp Figure S1**). The samples were then diluted to equal concentrations of 0.5 mg/ml using ultrapure water and *Rhodobacter* LPS was further buffer exchanged to ultrapure water using PES filter units with 5kDa cutoff (VivaSpin). Here, the sample was diluted with water 1:1 and concentrated to the original volume with the process repeated four times to remove small molecule carbohydrates. In case of OMVs isolated from *R. capsulatus* B10 and undiluted sample was used for the assay. For the phage sample, the ion exchange-purified phage was used in the assay. One *µ*l of the phage was added to 5 *µ*l of the sample (OMVs, LPS or ultrapure water) that was premixed with 50mM Tris, pH=7.5 to yield a final 10mM concentration of Tris. The phage-sample mixture was incubated 10 min at room temperature and then diluted 100 times in YPS. Serial dilutions were made and dropped onto plates with soft agar containing 75 *µ*l of overnight culture of SB1003. The plates were incubated overnight at 30°C and plaques were counted the next morning.

### Isolation, sequencing and analysis of the phage genome

The genomic ssDNA was isolated from the ion exchange-purified sample of the phage using a Virus DNA kit (Macherey-Nagel). For sequencing on the Oxford Nanopore platform, libraries were prepared using the SQK-RBK114-24 Rapid Sequencing kit (Oxford Nanopore Technologies) according to the manufacturer’s instructions. The libraries were sequenced with a FLO-FLG114 flow cell in a MinION device controlled by MinKNOW software v.23.07.12, (Oxford Nanopore Technologies), which was also used for basecalling (Super-accuracy model for genome assembly, and Super-accuracy model for modified base detection for 5mC, 5hmC, and 6mA in any contexts), demultiplexing and barcode trimming. Complete genome sequences were obtained using Flye v.2.9.1 (43) and Medaka consensus pipeline v1.7.2 (Oxford Nanopore). Methylation levels were analyzed by modkit v0.2.7 (Oxford Nanopore Technologies) using a 10% percentile threshold (A: 0.5996094, C: 0.60546875) for calls to pass. Only passed calls were included in the summary statistics. ORFs were identified using *ab initio* prediction of Prokka 1.14.5 (44) and GeneMarkS 4.28 (45) with the ‘phage’ algorithm. The annotation and circular maps were done in Artemis (46). The predicted function of protein was based on primary sequence similarity using blastp (47), and the comparison of profile hidden Markov models using HHpred (30). The prediction of protein structures was performed using AlphaFold2 (48).

### Phylogenetic analysis

For initial taxonomic placement, protein sequences of Ebor were used as input to MOP-UP (49), which was run with minimum protein identity cutoffs of 30% and 50%. To construct a phylogenetic tree of the *Microviridae*, representative VP1 (major capsid protein) sequences for each microviral ‘family’ as defined by Kirchberger *et al*. (4), and two additional sequences for phage Ebor and ‘Ascunsovirus oldenburgii’ ICBM5 (20), were chosen. Proteins were aligned using ClustalOmega using the mBed algorithm (50) for initial and refinement iteration guide trees (cluster size 100) and 100 refinement interactions. After pruning sites with > 50 % gaps in the alignment, a phylogenetic tree was constructed from the resulting alignment in RAxML (51) using the PROT_GAMMA_GTR substitution model and 100 rapid bootstrapping replicates. Transfer bootstrap estimates were then assessed using the BOOSTER web interface (52).

### Proteomic analysis

The LC-MS/MS analysis was performed by The Centre of Excellence in Mass Spectrometry, University of York. Here, the samples of purified Ebor were run 1 cm into a 10% SDS PAGE gel and stained with SafeBLUE stain (NBS Biologicals). The stained gel segment was excised, destained, reduced with DTE and alkylated with iodoacetamide. Protein was in-gel digested using Promega sequencing grade trypsin and incubated overnight at 37°C. Resulting peptides were analysed by LC-MS/MS over a 1 h acquisition with elution from a 50 cm EN C18 PepMap column (Thermo Fisher Scientific) driven by a mClass UPLC (Waters) onto an Orbitrap Fusion Tribrid mass spectrometer (Thermo Fisher Scientific) operated in DDA mode. MS1 spectra were acquired at high resolution in the Orbitrap mass analyser. MS2 spectra were acquired in TopSpeed mode in the linear ion trap after HCD fragmentation.

The LC-MS chromatograms were aligned using Progensis QI and a concatenated .mgf file searched against the *R. capsulatus* SB 1003 subset of NCBI and the microvirus proteome with all translated ORFs as identified by ORFfinder (NCBI), appended with common contaminants. Peptide identifications were filtered to 1% FDR using the Percolator algorithm before importing accepted identifications onto the chromatographic data in QI and matching identifications between runs. Data were further filtered to require a minimum of two unique peptides per protein. Quantification has been performed using a Top3 approach; taking the peak areas from the best three responding peptides as a proxy for each protein. The summed, integrated peak areas for the Top3 peptides are then converted to an estimation of relative molar percent within the sample by expressing each as a percentage of the sum total Top3 areas among all proteins.

### Production of the lytic enzyme and enzymatic assays

The *gp7* was cloned using HiFi assembly (New England Biolabs) of PCR-generated products with pETYSBLIC3c plasmid (53). The primers used for amplification were 5’ TCC AGG GAC CAG CAA TGA TCT ATC AAG GCA AAG ACC 3’ acting as a forward primer and 5’ TGA GGA GAA GGC GCG TCA AAG CCA GTC CAC CGA ATG 3’ acting as a reverse primer. The assembled plasmid was transformed into *E. coli* Stellar cells (Takara Bio) and individual colonies were verified for the presence of the insert by sequencing (Eurofins Genomics GATC). The plasmid was subsequently isolated from a verified colony and transformed to expression strain E. coli Rosetta (DE3) pLysS (Novagen). The expression was induced with 0.5 mM IPTG at 25°C. The pellet was resuspended in buffer Lys (20mM Tris, pH=7.5; 200mM NaCl; 5mM CaCl_2_; 20mM Imidazole), sonicated using Sonoplus HD2070 (Bandelin) and purified using Super Ni-NTA resin (Generon). The lysate was incubated with the resin for 15 minutes and subsequently washed three times with buffer Lys. Then, modified buffer Lys with an increased Imidazole concentration to 250mM was added to the resin, incubated for 5 min and the eluted sample was buffer exchanged using 10kDa filter units to the original buffer Lys. The his-tag was cleaved overnight in the presence of 1mM DTT and 20 mg of 3C protease. His-tag and the protease were removed from the protein samples by 20 min incubation with Super Ni-NTA resin (Generon). The sample was then loaded through a Superdex 75 10/300 column using the AKTA portal to separate the lysozyme from any excess protease, and the sample was then concentrated to 0.5 mg/ml using 10kDa Vivaspin^TM^ (Cytiva) filter units. Zymogram and plate lysis assays were adapted from Benesik et al., 2018 (54) and performed as described previously (3).

### Cryo-EM and ET sample preparation and data acquisition

Four µl of purified virus sample were applied onto glow-discharged Quantifoil^®^ R2/1 300 mesh copper grids and blotted using Vitrobot Mark IV (Thermo Fisher Scientific) for 2 sec, blot force 0, wait time 8 sec, and plunge frozen in liquid ethane. For the virus attached to cells, cells of the respective strain were grown until the stationary phase in RCV containing a quorum-sensing molecule O-acetylserine (4 mM). Then, 1.5 ml of the culture was spun at 1000 × g/5 min, the supernatant spun at 14 000 × g/15 min and the final pellet resuspended in 75 µl of RCV. Then, 1 µl of ion exchange-purified phage was added to 4 *µ*l of cells and after 5 min of incubation mixed 1:2 (v/v) with 6 nm BSA-gold tracers (Aurion) which were transferred to RCV by Micro Bio-Spin® 6 columns (Biorad). Four *µ*l of the mixture were applied onto glow-discharged Quantifoil^®^ R2/1 200 mesh copper grids. The grids were blotted from the back side by covering one pad of Vitrobot with parafilm. Vitrobot settings of blotting time 9 sec, blot force 0, wait time 2 sec, were used before plunge freezing in liquid ethane. A similar approach was used for freezing Ebor with LPS and OMVs, with the blotting time decreased to 5 seconds. All grids were screened using a Glacios TEM (Thermo Fisher Scientific) operated at 200 kV and equipped with Falcon 4 camera at the cryo-EM facility of the University of York. Automatic data acquisition was performed using microscopes at York, Eindhoven and eBIC (Harwell) as stated in **Supp Table S1**. The raw data was acquired in EER format for Falcon 4(i) and TIFF format for K3, and motion corrected using RELION implementation of MotionCor2 (55,56). The dose-weighting was performed using the same implementation in the case of single particle data and using the alignframes IMOD command (57) in the case of tilt series. This command was also used for generating stacks out of individual tilt images. For single particle data, the CTF estimation was performed using CTFFIND4 (58).

### Cryo-EM single particle analysis of the empty particle dataset

The particles were picked using a template-based autopicking pipeline implemented in Relion 3.1 (56). For the template generation, 200 particles were manually picked, extracted and 2D classified. Resulting class averages were used as 2D templates for autopicking, identifying 3976 particles. The particles were extracted in an unbinned 256 px box and were subject to further 2D classification to discard bad particles. The subset of 2880 particles was selected for further processing with applied I4 symmetry. The initial model was generated *de novo* using Relion 3.1. The 3D refinement was performed followed by 3D classification to discard bad particles. Then, a mask was generated using the volume segmentation tool in UCSF ChimeraX (59) and relion_mask_create command, which encompassed the density of the capsid shell. After another run of 3D refinement and classification, the final 1984 particles were Bayesian-polished and CTF refined using Relion 3.1. The final map was improved using DeepEMhancer (60).

### Cryo-EM single particle analysis of the native particle dataset

The particles were picked using a template-based autopicking pipeline implemented in Relion 3.1 (56). For the template, the 3D map obtained from the empty particle dataset was used. The autopicking identified 5271 particles, which were extracted in an unbinned 360 px box, and subjected to further 2D classification to discard bad particles. The subset of 5043 particles was selected for further processing with applied I4 symmetry. The map generated from the empty dataset was low-pass filtered to 45 Å and used as an initial model. The 3D refinement was performed followed by 3D classification to discard bad particles. Then, a mask was generated using the volume segmentation tool in UCSF ChimeraX (59) and the relion_mask_create command, which encompassed the density of the capsid shell. After another run of 3D refinement and classification, the final 4935 particles were Bayesian-polished and CTF refined using Relion 3.1 (56,61). The final map was improved using DeepEMhancer (60).

### Cryo-EM single particle analysis of the genome-releasing particle dataset

The particles were manually picked to begin training the Topaz software for autopicking. The model was then used to autopick 4891 particles. The particles were extracted in an unbinned 360 px box, rescaled to 64 px, and subjected to 2D classification. Selected classes were refined in C5 symmetry against a *de novo* model generated using Relion 3.1 (56). One round of 3D classification was applied to further separate particles that lost the capsomer and autorefined (56).

### Model building

The protein model was first predicted using AlphaFold2 (48) and fitted into the deepEMhanced map using UCSF ChimeraX 1.7.1 (59). The model was refined iteratively using real space fitting in Coot 0.9.8.5 (62) and real space refinement in Phenix 1.20.1 (63). Models were subsequently refined using NCS constraints and with the interacting partners in the virion, to prevent inter-molecular atom clashes. During the iterative refinement process, the molecular geometry was monitored using the wwPDB validation server [https://validate.wwpdb.org] and MolProbity (64), suspicious outliers were fixed manually using Coot and further refined in Phenix. The refinement statistics are presented in **Supp Table S4**.

### Tomogram generation and subtomogram averaging

The tomogram generation, CTF estimation and subtomogram averaging were performed in EMAN2 (65). The particles were picked using reference-based picking and manually corrected to retain only particles attached to the sample and remove those present on carbon. The subtomogram averaging was performed as shown in **Supp Figures S5**-**6**. After the final reconstruction with icosahedral symmetry applied, the information about subtilt alignment was utilized to polish the tomograms using the e2spt_polishtomo.py command of EMAN2 (65). The final maps from classification were filtered using e2proc3d.py command according to their respective refinement map to minimize the low-resolution ice gradient effect on their density.

### Cryo-EM single particle analysis of the particles attached to cells

The particles were picked using Topaz (66) with the neural network trained on a set of particles manually picked from 25 random micrographs. The figure of merit for particle extraction was selected based on the manual inspection of 25 micrographs, the particles were extracted in an unbinned 256 px box and processed in RELION 5 (67) as shown in **Supp Figure S7**. The final maps were low pass filtered to their estimated maximum resolution using relion_image_handler command.

## Data visualization

The structural figures were created using UCSF ChimeraX 1.7.1 (59). The tomograms were visualized and movies were made using IMOD 4.12 (57). The charts were plotted using the ggplot2 system of the package R [https://www.r-project.org/]. The electrostatic potential was coloured according to Adaptive Poisson-Boltzmann Solver (68). The segmentation of tomograms was done using EMAN2 (69).

## Acknowledgments

The research leading to these results has received funding from the Wellcome Trust grants 224067/Z/21/Z to P.B., 206377 to A.A.A., 109363/Z/ 15/A to P.F; a grant (RGPIN 2018-03898) from the Canadian Natural Sciences and Engineering Research Council (NSERC) to J.T.B.; N.I.H. Grant 1P20GM152333-01:8040 to P.C.K.; and the German Academic Exchange Service (DAAD) and MITACS (project ID 28535) to H.C.K. This work was supported by a National Institute of Virology and Bacteriology project (Program EXCELES, ID project no. LX22NPO5103) funded by the European Union - Next Generation EU. The imaging was done at the University of York cryo-EM facility supported by the Wellcome Trust (206161/Z/17/Z). We greatly acknowledge Diamond for access and support of the cryo-EM facilities at the UK national electron Bio-Imaging Centre (eBIC), proposal BI34172. We thank Thermo Fisher Scientific and Dr Ieva Drulyte for outstanding tilt series acquisition training, and Dr Oliver Raschdorf for the data acquisition assistance. Further tomogram acquisition and processing training was obtained at the Stanford-SLAC Cryo-EM Center (S2C2), which is supported by the National Institutes of Health Common Fund Transformative High-Resolution Cryo-Electron Microscopy program (U24 GM129541) and the invaluable support of Dr Yan Liu and Prof Wah Chiu. We also thank Dr Muyuan Chen (Stanford-SLAC), Dr Dominik Hrebik (Max Planck Institute of Biochemistry), Jake Smith (Rosalind Franklin Institute), and Prof Bentley Fane (University of Arizona) for fruitful discussions and Prof Alan Davidson (University of Toronto) for mentoring P.B.

The Viking cluster was used during this project, which is a high-performance compute facility provided by the University of York. We are grateful for computational support from the University of York, IT Services and the Research IT team. Further computing was performed on workstations purchased thanks to financial support from the Hartshorn-Jones fund at the University of York. Mass spectrometry analysis was performed at the York Centre of Excellence in Mass Spectrometry supported by a major capital investment through Science City York and Yorkshire Forward with funds from the Northern Way Initiative and EPSRC (EP/K039660/1; EP/M028127/1). We thank Vytaute Jurgeleviciute (University of York) for proof-reading.

## Author contributions

P.B., conceptualization (equal), data curation (lead), formal analysis (equal), funding acquisition (equal), investigation (lead), methodology (lead), project administration (equal), supervision (supporting), validation (lead), visualization (lead), writing - original draft (equal), writing – review and editing (equal); C.I.W.M., conceptualization (supporting), data curation (supporting), formal analysis (supporting), investigation (supporting), methodology (supporting), writing - original draft (equal), writing – review and editing (supporting); P.C.K., conceptualization (supporting), data curation (supporting), formal analysis (supporting), funding acquisition (supporting), investigation (supporting), methodology (supporting), writing - original draft (supporting), writing – review and editing (supporting); H.T.J., formal analysis (supporting), methodology (supporting), writing – review and editing (supporting); T.B., data curation (supporting), formal analysis (supporting), funding acquisition (supporting), investigation (supporting), methodology (supporting), writing – review and editing (supporting); L.B., data curation (supporting), formal analysis (supporting), investigation (supporting); N.T.B.A., formal analysis (supporting), investigation (supporting), methodology (supporting),; D.A.K.T., data curation (supporting), investigation (supporting), methodology (supporting); H.C.K., investigation (supporting); T.R.N., investigation (supporting), methodology (supporting); M.C., supervision (supporting); S.J.H., investigation (supporting); J.P.T., investigation (supporting); J.N.B., funding acquisition (supporting), methodology (supporting), supervision (supporting), writing – review and editing (supporting); J.T.B., formal analysis (supporting), funding acquisition (supporting), investigation (supporting), methodology (supporting), supervision (supporting), writing – review and editing (supporting); P.C.M.F., conceptualization (supporting), formal analysis (equal), funding acquisition (equal), investigation (supporting), methodology (supporting), project administration (supporting), supervision (supporting), writing - original draft (supporting), writing – review and editing (equal); A.A.A, conceptualization (equal), formal analysis (equal), funding acquisition (equal), methodology (supporting), project administration (equal), supervision (lead), writing - original draft (supporting), writing – review and editing (equal)

## Data availability

Ebor nucleotide sequence data are available in DDBJ/ENA/GenBank: PP754850.1. Raw LC-MS/MS data of purified Ebor are deposited at MassIVE: MSV000094685. Cryo-EM electron density maps have been deposited in the Electron Microscopy Data Bank, https://www.ebi.ac.uk/pdbe/emdb/ and the fitted coordinates have been deposited in the Protein Data Bank, www.pdb.org with all the codes listed in **Supp Table S1**. The authors declare that all other data supporting the findings of this study are available within the article and its Supplementary Information files or are available from the authors upon request.

