## Supplementary information for "A stargate mechanism of *Microviridae* genome delivery unveiled by cryogenic electron tomography"

Pavol Bardy *et al.*

### Supplementary information

**Description of Supp Data**
**Supp Figures S1-S8**
**Supp Tables S1-S4**
**Supplementary References**

### Description of Supp Data

**Supp Movie S1:** Tomogram of Ebor particles attached to outer membrane vesicles formed from disrupted cells of *Rhodobacter capsulatus*, strain B10.

**Supp Movie S2:** Tomogram of Ebor particles attached to the host cell of *Rhodobacter capsulatus*, strain B10.

**Supp Movie S3:** Depiction of the stargate opening of Ebor particle, side view.

**Supp Movie S4:** Depiction of the stargate opening of Ebor particle, top view.

**Supp Data S1:** Mass spectrometry analysis of purified virions of Ebor.

**Supp Data S2:** Database of *Microviridae* genomes used for the phylogeny analysis.

**Supp Data S3:** A model of the trimeric protrusion predicted by AlphaFold2.

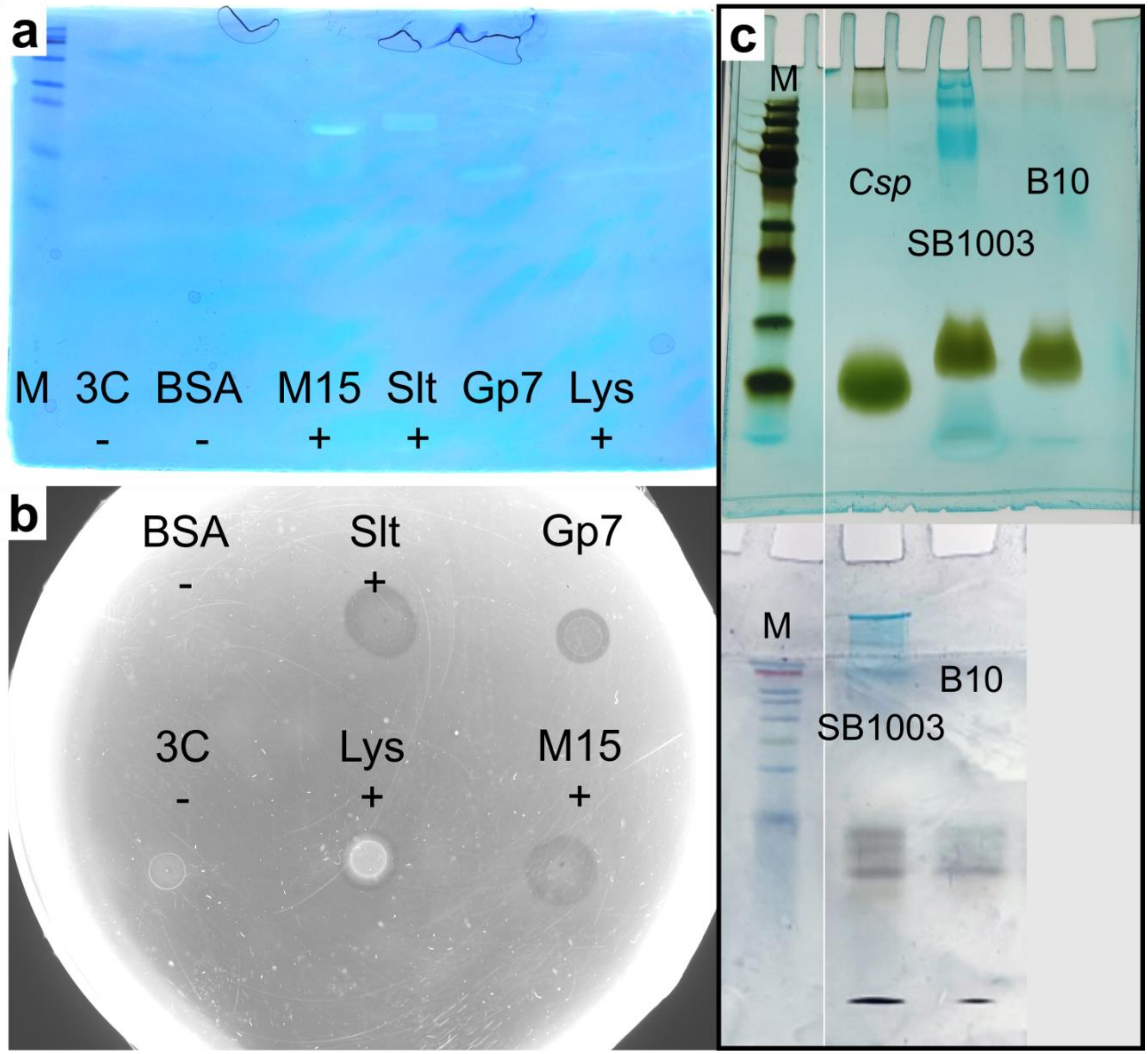


**Supp Figure S1: Images of gels and the water agar plate.** a-b) Enzymatic assays of Gp7 with zymogram SDS-PAGE gel (a) and water agar plate (b) shown. BSA and 3C protease were used as negative controls, M15 and Slt enzymes of *Rhodobacter* phage Jorvik (1) as well as lysozyme were used as positive controls. c) Gels with LPS samples that were used for Ebor inhibition assay ran on Any kD™ Mini-PROTEAN® TGX™ gradient gel (top) and more diluted samples ran on a 15 % polyacrylamide gel (bottom).

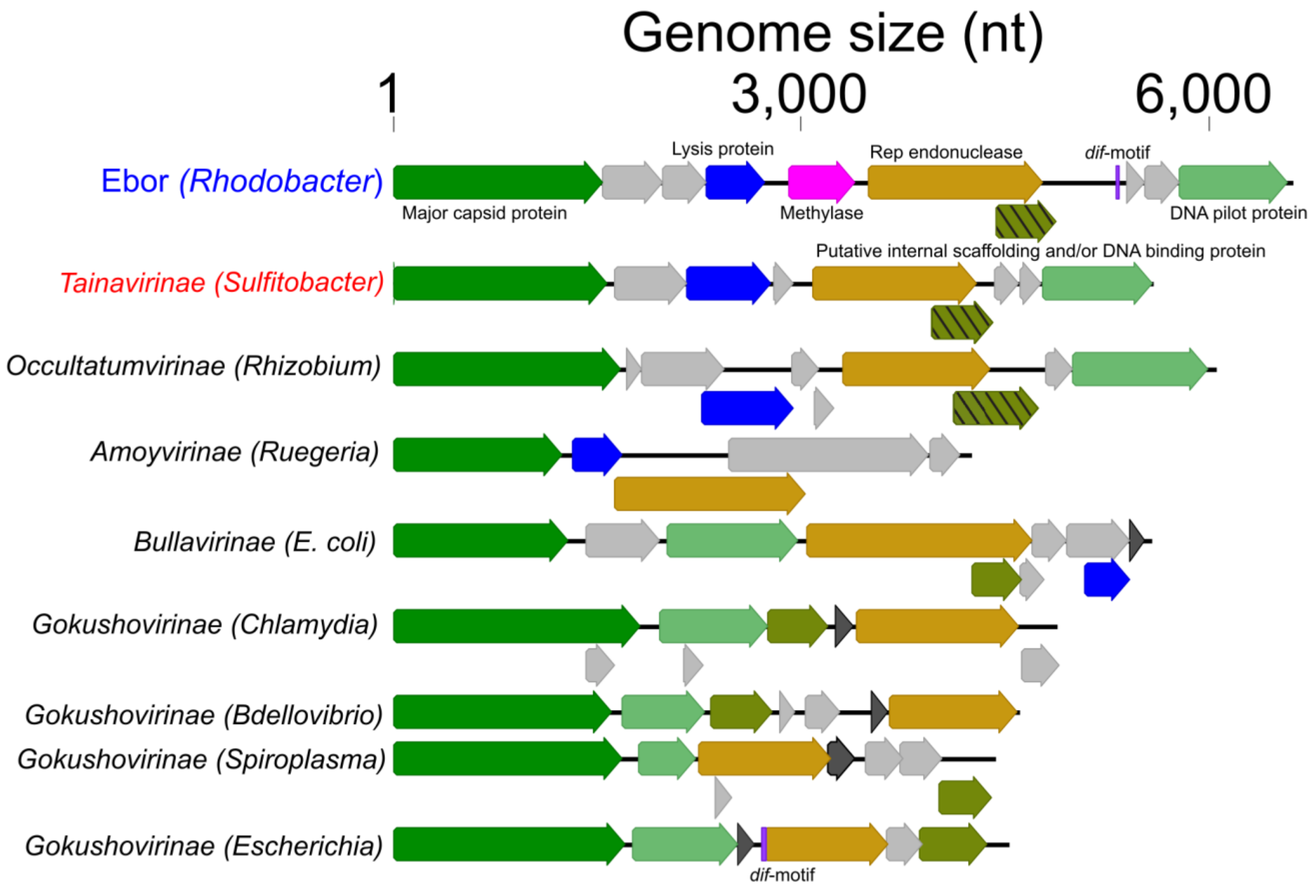


**Supp Figure S2: Genome structure of Ebor and select other microviruses.** Homologous/analogous proteins are shown in the same colour.

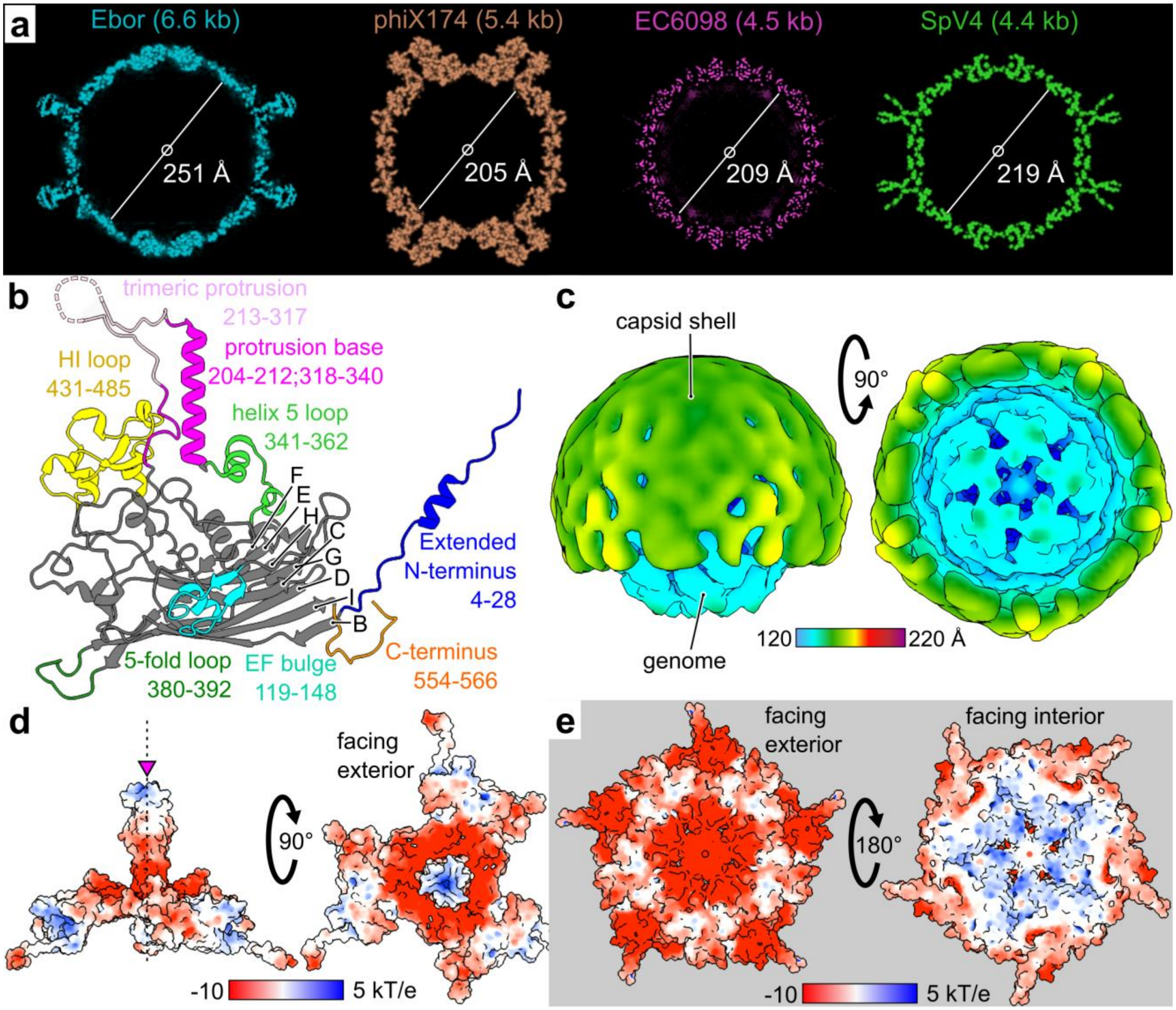


**Supp Figure S3: Additional structural analysis of the native virion of Ebor.** a) Diameters of individual *Microviridae* phages. Central Z slices of the capsids are shown, the densities of phiX174 and SpV4 were generated by ChimeraX molmap command (2) of PDB bioassemblies 2BPA_A and 1KVP_A respectively. For EC6098, a segmentation of EMD-27397 was used, masking out the central virion density for clarity. b) Ribbon diagrams of the major capsid protein of phage EC6098, PDB code 8DES_A. Regions of interest are highlighted in colour and delimiting residues numbers are shown. c) Map of Ebor virion releasing the genome *in vitro* reconstructed by single particle analysis of variant S120 virions purified using CsCl gradient. d-e) electrostatic potential of Ebor protrusion (d) and penton (e) estimated according to Adaptive Poisson-Boltzmann Solver (3). The AlphaFold2-predicted (4) protrusion loops were connected into a single model with respective subunits of major capsid protein in Coot (5) with the axis of symmetry highlighted (magenta triangle). The colour coding corresponds to the electrostatic potential values of the protein surface calculated at T=298.15K and pH=7.0.

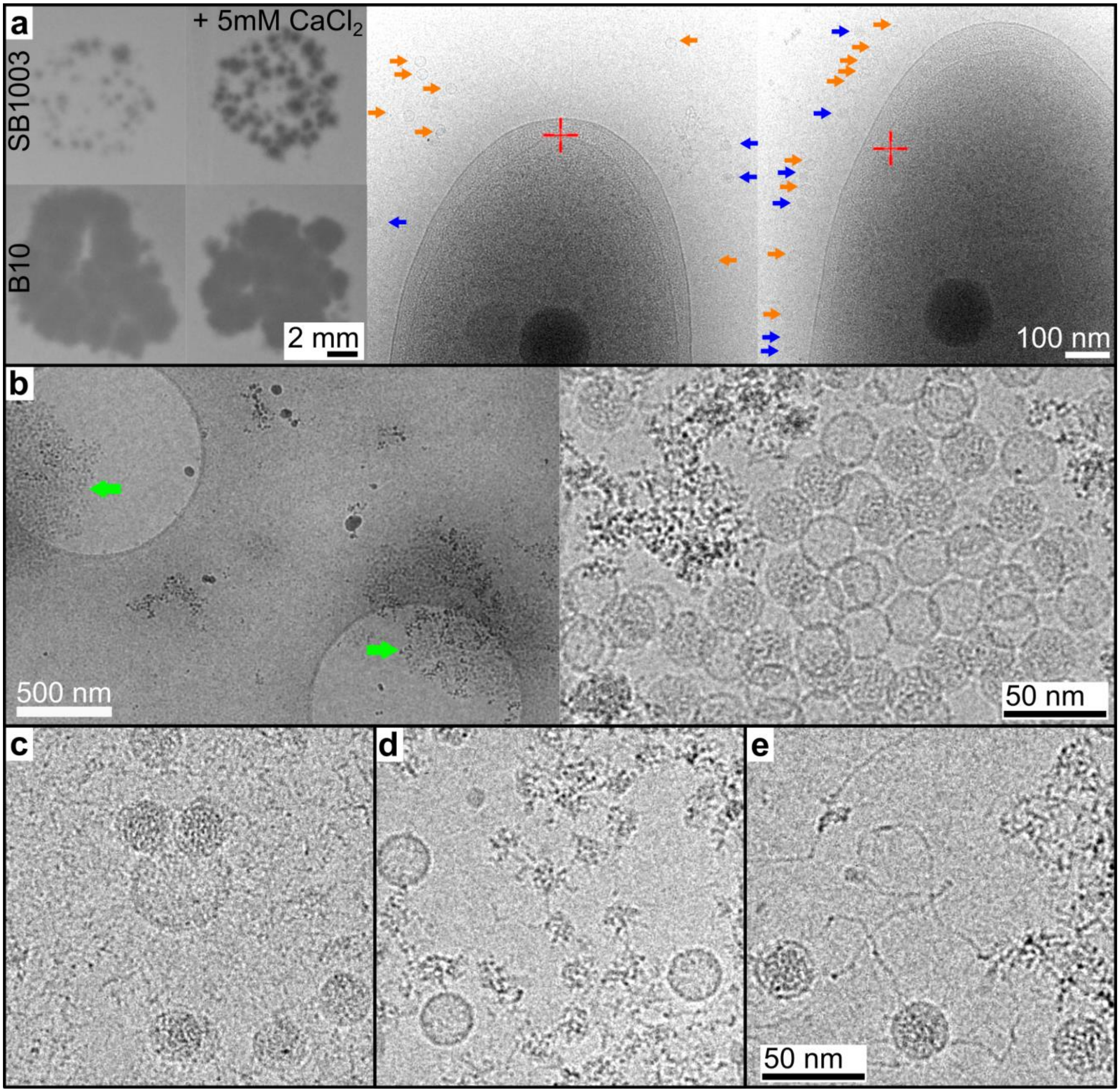


**Supp Figure S4: Additional cryo-EM images of Ebor.** a) The same virus stock as used in the experiments shown in the **Figure 3d-f)** was imaged after 5 min of incubation with the host strain SB1003 to which 5 mM $CaCl_2$ was added. The effect of the calcium addition on the plaque size is shown in left. b-e) Images of Ebor particles purified using different methods. Ebor variant R120 purified using ion exchange chromatography (b). The particles formed aggregates (green arrows) and the ratio of empty and native particles was close to 1:1. Ebor variant R120 purified using CsCl ultracentrifugation (c), all observed particles were native. Ebor variant S120 purified using sucrose ultracentrifugation (d), all observed particles were empty, the contaminants are likely ejected genome molecules. Ebor variant S120 purified using CsCl ultracentrifugation (e), the particles appear to be releasing their genome. Images were collected at 200 kV using a Glacios TEM equipped with Falcon 4 camera, with the total exposure dose of ~3 $e^{-}/Å^{2}$ in case of cells and 50 $e^{-}/Å^{2}$ in case of purified viruses.

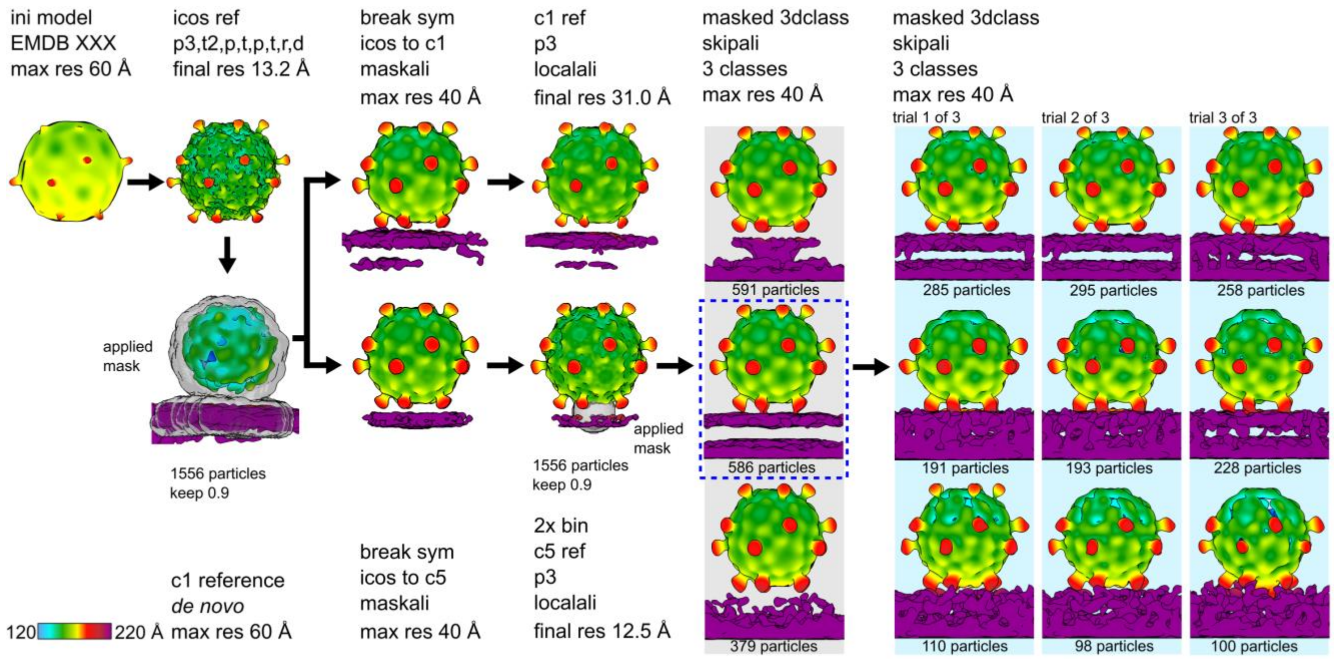


**Supp Figure S5: EMAN2 pipeline (6) applied for the subtomogram reconstruction of the dataset of phage Ebor attached to cells.** The colour bar indicates the distance from the centre of the capsid. Different iterations of the refinement algorithm are labelled according EMAN2 conventions: p, 3D particle orientation; t, 2D subtilt translation; r, subtilt translation and rotation; d, subtilt defocus refinement.

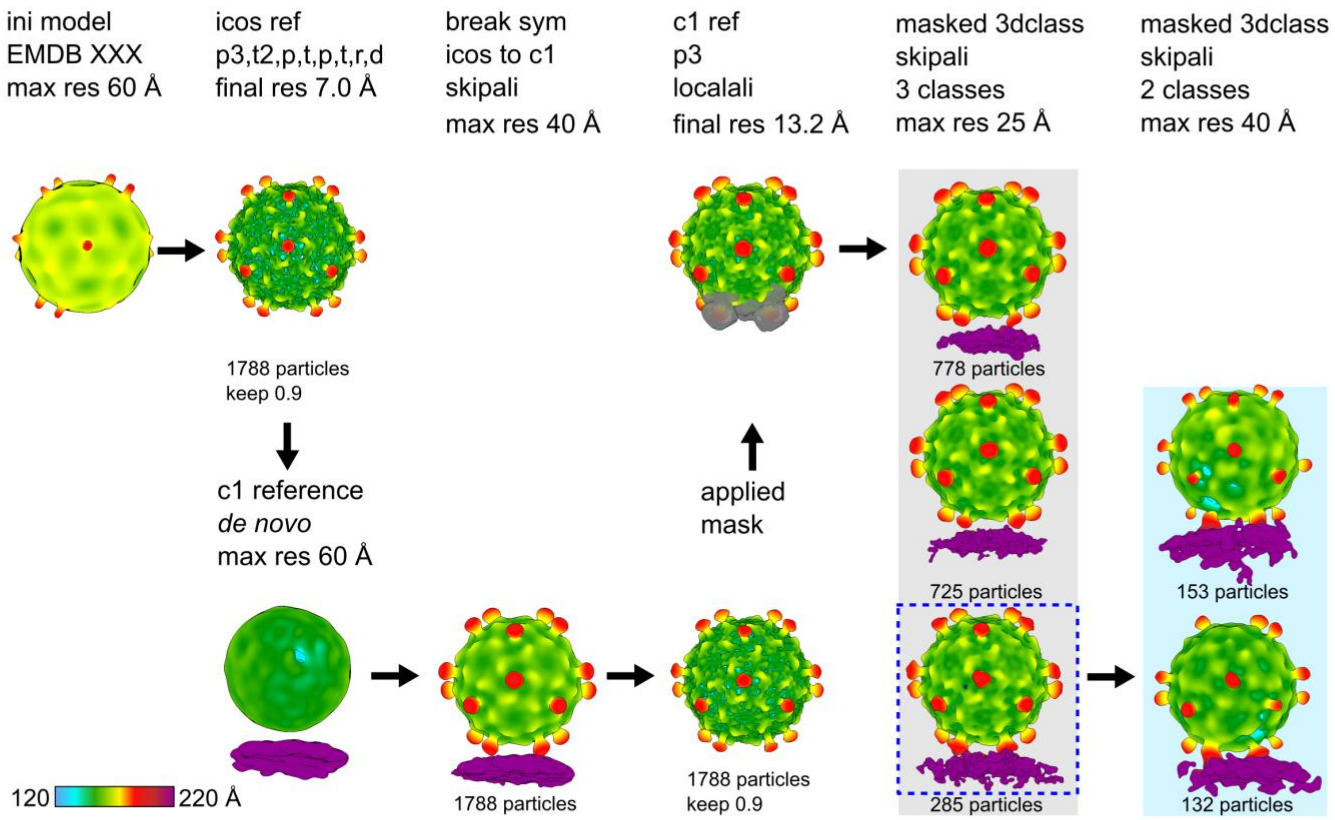


**Supp Figure S6: EMAN2 pipeline (6) applied for the subtomogram reconstruction of the dataset of phage Ebor attached to OMVs.** The colour bar indicates the distance from the centre of the capsid. Different iterations of the refinement algorithm are labelled according EMAN2 conventions: p, 3D particle orientation; t, 2D subtilt translation; r, subtilt translation and rotation; d, subtilt defocus refinement.

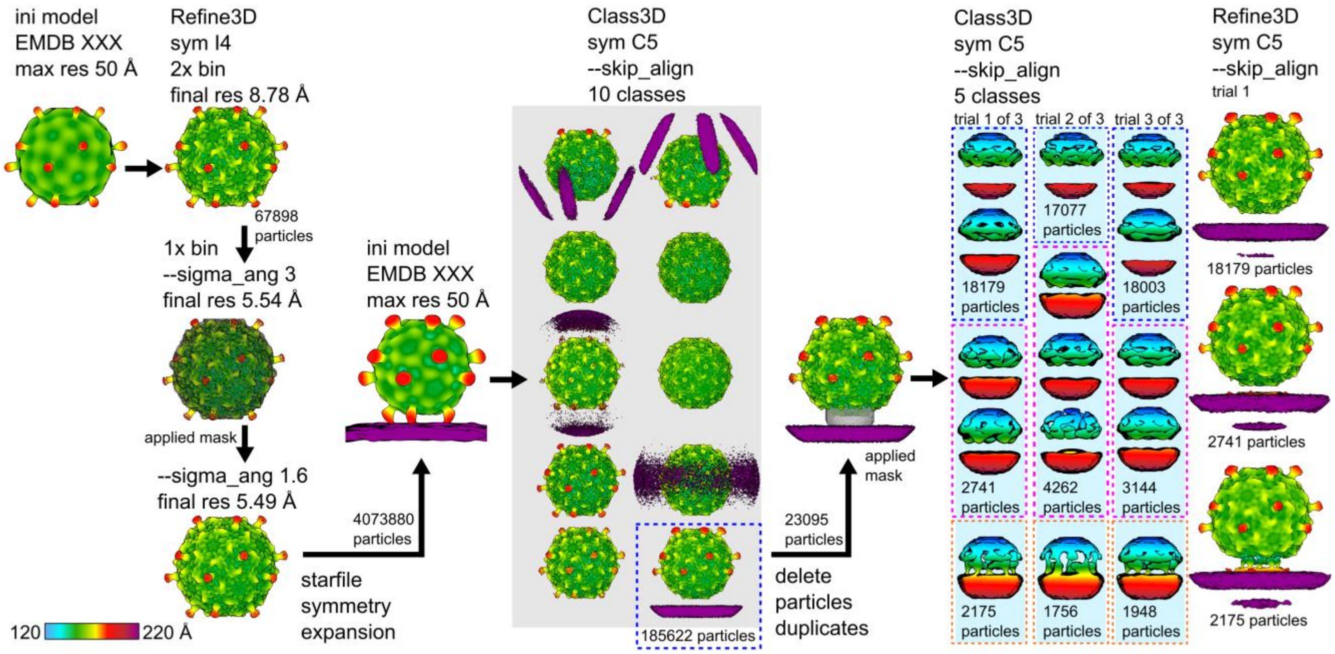


**Supp Figure S7: RELION pipeline** (7–9) **applied for the single particle reconstruction of the dataset of phage Ebor attached to cells.** The colour bar indicates the distance from the centre of the capsid.

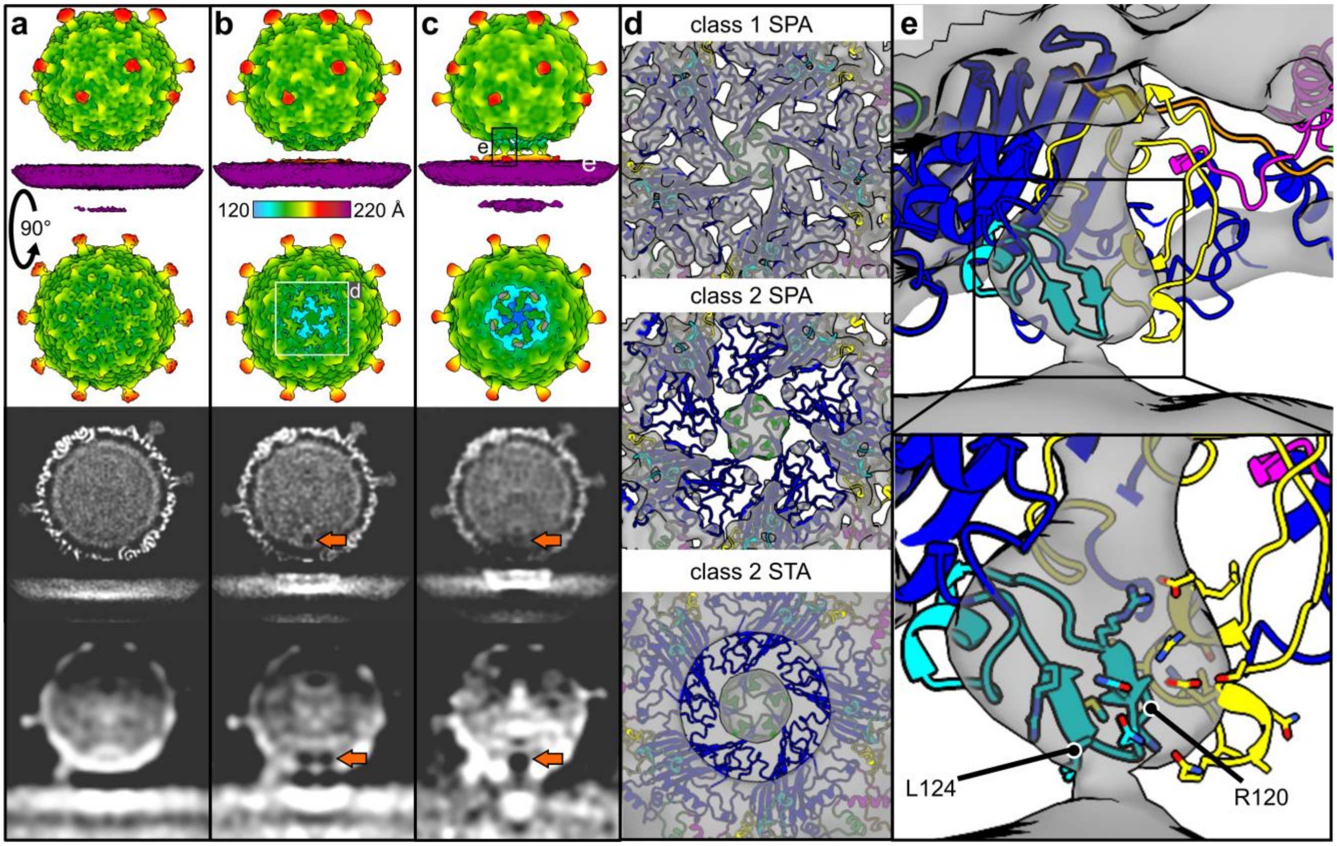


**Supp Figure S8: Single particle analysis of Ebor attached to cells.** a-c) C5 symmetry maps of the attached particles classified into class 1 (a), 2 (b) and 3 (c) reconstructed by single particle analysis (SPA). Side view, top view and central slice through y axis are shown from top to bottom, with a central slice through y axis of unfiltered maps reconstructed by subtomogram averaging (STA) shown on the very bottom for comparison. The colour bar indicates the distance from the centre of the capsid. The arrows point at the density of an internal cavity formed above the interacting penton. d) Fitting of the major capsid protein model shown as ribbon diagrams into the interacting penton of classes 1 and 2 reconstructed by SPA s and class 2 reconstructed by STA. A similar pore was identified in class 2 reconstructed by both methods, with detached density present right on the 5-fold axis. e) Fitting of the major capsid protein in class 3 map reconstrcuted from SPA shows the density connecting the capsid with the membrane corresponds to the position of EF bulge. Close up of the region is shown in the bottom rectangle. Individual residues are shown as sticks with R120 and L124 highlighted.

**Supp Table S1: Cryo-EM and ET data collection and processing information.**

| Parameters | Single particle analysis | | | | Subtomogram averaging | |
|---|---|---|---|---|---|---|
| | native particle | empty particle | genome-releasing particle | particles attached to cells | particles attached to cells | particles attached to OMVs |
| **Data collection** | | | | | | |
| Magnification | x120000 | x120000 | x120000 | x42000 | x45000 | x45000 |
| Pixel size [Å] | 1.20 | 1.20 | 1.20 | 2.143 | 2.50 | 3.15 |
| Camera (energy filter) [manufacturer] | Falcon 4 [TFS] | Falcon 4 [TFS] | Falcon 4 [TFS] | K3 (GIF) [Gatan] | Falcon 4i (Selectris) [TFS] | Falcon 4 [TFS] |
| Instrument* | Glacios, York | Glacios, York | Glacios, York | Krios II, eBIC | Glacios 2, TFS facility, Eindhoven | Glacios, York |
| Voltage [kV] | 200 | 200 | 200 | 300 | 200 | 200 |
| Data collection software | EPU (holey carbon set up) | EPU (holey carbon set up) | EPU (holey carbon set up) | EPU (lacey carbon set up) | Tomo5 | Tomo5 |
| Total exposure dose [$e^-/Å^2$] | 50 | 50 | 50 | 27.878 | 117.66 | 115.9 |
| Exposure dose per tilt [$e^-/Å^2$] | 50 | 50 | 50 | 27.878 | 3.18 | 1.9 |
| Number of tilts | 1 | 1 | 1 | 1 | 37 | 61 |
| Tilt span [°] | 0 | 0 | 0 | 0 | ±54 | ±60 |
| Tilt increment [°] | NA | NA | NA | NA | 3 | 2 |
| Number of acquired micrographs/tilt series | 996 | 1099 | 2810 | 2750 | 42 | 71 |
| Number of used micrographs/tilt series | 954 | 764 | 2095 | 2742 | 27 | 25 |
| Defocus range [µm] | 0.5 - 2.5 | 0.5 - 2.5 | 0.5 - 2.5 | 0.75 - 3.0 | 1.5 - 4.0 | 1.0 - 3.0 |
| **Data processing** | | | | | | |
| Reconstruciton software | RELION3 | RELION3 | RELION3 | RELION5 | EMAN2 | EMAN2 |
| Symmetry | i4 | i4 | C5 | i4, symmetry expansion, c5 | icos, symmetry break to c1 and c5 | icos, symmetry break to c1 |
| Initial number of particles | 5271 | 3976 | 6302 | 75291 | 1593 | 1788 |
| Final number of particles | 4935 | 1984 | 401 | class1=18179, class2=2741, class3=2175 | c1=1556, class1=295, class2=193, class3=198 | c1=1788, weak1=725, weak2=778, strong1=153, strong2=132 |
| Initial model | emd-50356 | *de novo* | *de novo* | emd-50361 | emd-50357, break to c1 *de novo* | emd-50357, break to c1 *de novo* |
| Map resolution [Å] | 3.2 | 3.3 | 24 | 7.8, 10.4, 13.4 | c1=31, far=40†, close=40†, fused=40† | c1=13.2, weak1=25†, weak2=25†, strong1=40†, strong2=40† |
| FSC threshold | 0.143 | 0.143 | 0.143 | 0.143 | 0.2 | 0.2 |
| **Database entry** | | | | | | |
| EMDB | 50357 | 50356 | 50358 | 50359 | 50361 | 50360 |
| PDB | 9FFH | 9FFG | NA | NA | NA | NA |

**all manufactured by TFS; †low passed to this resolution; OMV, outer membrane vesicle; TFS, Thermo Fisher Scientific*

**Supp Table S2:** Raw data related to the Ebor inhibition assay.

| **Sample** | **OMVs** | **LPS *C.sph.*** | **LPS B10Δ*gtaI*** | **LPS SB1003Δ*gtaI*** |
|---|---|---|---|---|
| biol. rep. 1 buffer [PFUs] | 2.00E+10 | 3.20E+08 | 3.20E+07 | 6.00E+07 |
| biol. rep. 1 sample [PFUs] | 5.00E+08 | 3.00E+08 | 1.50E+06 | 1.10E+07 |
| biol. rep. 2 buffer [PFUs] | 2.00E+10 | 3.20E+08 | 6.00E+07 | 6.00E+07 |
| biol. rep. 2 sample [PFUs] | 5.30E+08 | 2.30E+08 | 1.70E+07 | 1.20E+07 |
| biol. rep. 3 buffer [PFUs] | 2.50E+09 | 3.20E+08 | 6.00E+07 | 6.00E+07 |
| biol. rep. 3 sample [PFUs] | 3.00E+07 | 3.50E+08 | 1.80E+07 | 8.00E+06 |
| biol. rep. 1 RI [%] | 2.50% | 93.75% | 4.69% | 18.33% |
| biol. rep. 2 RI [%] | 2.65% | 71.88% | 28.33% | 20.00% |
| biol. rep. 3 RI [%] | 1.20% | 109.38% | 30.00% | 13.33% |
| **average RI [%]** | **2.12%** | **91.67%** | **21.01%** | **17.22%** |

*OMV, outer membrane vesicle; LPS, lipopolysaccharide; biol. rep., biological replicate; PFU, plaque-forming unit; RI, relative infection*

**Supp Table S3: Bacterial strains used in this study.**

| Bacterial species | strain | Purpose | Reference/source |
|---|---|---|---|
| *Rhodobacter capsulatus* | DE442 | source of Ebor | (10) |
| | SB1003 | propagation of Ebor R120, plating strain for inhibition assay | (11) |
| | SB1003Δ*gtaI* | capsuleless strain, LPS extraction | (12) |
| | B10 | propagation of Ebor S120, host for cryo-EM experiments | (1) |
| | B10Δ*gtaI* | capsuleless strain, LPS extraction | (13) |
| *Escherichia coli* | Stellar | cloning of gp7 | TakaraBio |
| | BL21(DE3) | expression of gp7 | Novagen |

**Supp Table S4: Model refinement statistics**. The values were calculated using wwPDB EM Validation server and Molprobity server (14).

| Parameters | | Model | |
|---|---|---|---|
| | | 9FFH | 9FFG |
| **Map** | EMDB code | 50357 | 50356 |
| | Resolution [Å] | 3.2 | 3.3 |
| **Model composition** | Non-hydrogen atoms | 3473 | 3379 |
| | Protein residues | 451 | 437 |
| **R.M.S. deviations** | Bond lengths [Å] | 0.26 | 0.24 |
| | Bond angles [°] | 0.55 | 0.50 |
| **Ramachandran plot** | Favoured [%] | 96.42 | 96.07 |
| | Outlier [%] | 0.45 | 0.00 |
| | Z-score | -1.06 ± 0.39 | 0.08 ± 0.42 |
| **Validation** | Rotamer outliers [%] | 1.83 | 1.07 |
| | Clashscore | 5.05 | 3.11 |
| | Molprobity score | 1.70 | 1.39 |
| | CaBLAM outliers [%] | 3.20 | 3.00 |
| **Atom inclusion*** | | 0.85 | 0.84 |

**at the recommended contour level = 0.1*
